# A wireless modular platform for neuro-behavioral recording and closed-loop manipulation in small animals

**DOI:** 10.64898/2026.08.25.747153

**Authors:** Zifang Zhao, Hongyu Chang, Praveen Paudel, Jaehyo Park, Can Liu, Maria Q. Aurelio, Azahara Oliva, Antonio Fernandez-Ruiz

## Abstract

Investigating the neural mechanisms of social group interactions and other naturalistic behaviors in small animals remains limited by current technology. Tethered neural recording systems are incompatible with many of these behaviors, while existing wireless devices for small animals are constrained by weight, bandwidth, recording duration, and the lack of closed-loop modulation capabilities. To overcome these limitations, we developed a Wireless, Interactive, Lightweight Datalogger (WILD) with integrated flexible neural probes, optogenetics, an inertial measurement unit, an ultrasonic microphone, and a head-mounted camera. This platform enables simultaneous, long-term recording of neural activity, locomotor variables, vocalizations, and eye movements from groups of freely moving mice in both laboratory and outdoor settings. Model-based real-time signal processing detects specific neural events and behavioral motifs to trigger closed-loop neural interventions. By combining multimodal recordings with advanced onboard signal-processing capabilities in a compact device, WILD enables the investigation of neural mechanisms underlying a broad range of natural behaviors in small animals.

## Introduction

A fundamental goal in neuroscience is to uncover the neural mechanisms of behavior. To achieve this goal, it is necessary to both record and precisely manipulate the activity of populations of neurons and relate it to quantitative descriptions of behavioral dynamics. Remarkable progress has been made in recent years in the development of tools for large-scale neural recordings^1,2^, manipulating the activity of specific cell types with high spatio-temporal resolution^3,4^ and accurately tracking behavioral and physiological variables^5–7^, particularly in laboratory rodents. However, due to technical limitations, these methods have been mostly deployed in reduced and simplified behavioral paradigms that fail to fully capture the richness and complexity of natural behaviors. As a result, there is an increasing need to expand the behavioral repertoire of mechanistic neuroscience studies to include more naturalistic and ecologically relevant behaviors^8,9^ such as social group dynamics, navigation, and foraging in large and complex environments, and even going beyond the laboratory to outdoor settings^10–12^. This need has led to a growing demand for approaches that enable mechanistic studies of neural activity in the context of unrestricted, naturalistic behaviors.

The most common methods for recording neural activity in freely behaving animals are the use of tethered electrophysiological or imaging devices. These approaches enable the simultaneous recording of large numbers of neurons, but at the cost of restricting animals’ natural movements, the size and complexity of the behavioral apparatuses, and the range of behaviors (e.g., group interactions). On the other hand, wireless recording methods enable neural recordings during unrestrained behavior, but they currently have substantial limitations. Battery weight and energy efficiency limit recording duration and electrode numbers, especially in small animals such as mice. Existing wireless neural recording interfaces are designed without considering the need for broader functionality, such as manipulating neural activity or incorporating other types of sensors to record behavioral and physiological variables^13–16^. Another key limitation of current wireless neural systems is their inability to perform complex on-board signal processing, which limits their application in studies requiring closed-loop neural intervention. Recent advances in machine learning (ML), particularly in developing lightweight model frameworks on embedded processors, offer a pathway towards overcoming this limitation^17^. Embedding efficient ML algorithms and artificial neural networks (ANN) could, in principle, endow wireless neural interfaces with advanced capabilities for real-time detection and manipulation of neural or behavioral events, but this application is yet to be developed.

To enable mechanistic studies of fully unrestricted natural behaviors in small animals, we developed WILD, an integrated platform for wireless recording of neural activity and multiple behavioral and physiological variables (Fig. 1a). Embedded real-time signal processing and efficient ANN architectures allow for wireless closed-loop optogenetic manipulations based on real-time detection of neural and behavioral events (Fig. 1a). We demonstrate the capabilities of this system in sets of experiments not feasible with currently existing approaches, including recordings of neural activity and behavioral variables (such as vocalizations, pupil dynamics and locomotion) during mice social group behavior, and neural and behaviorally triggered closed-loop manipulations in mice navigating complex 3D environments. To facilitate the adoption of this system across different laboratories, all hardware and software elements of this platform are open-source and fully documented.

**Figure 1:**
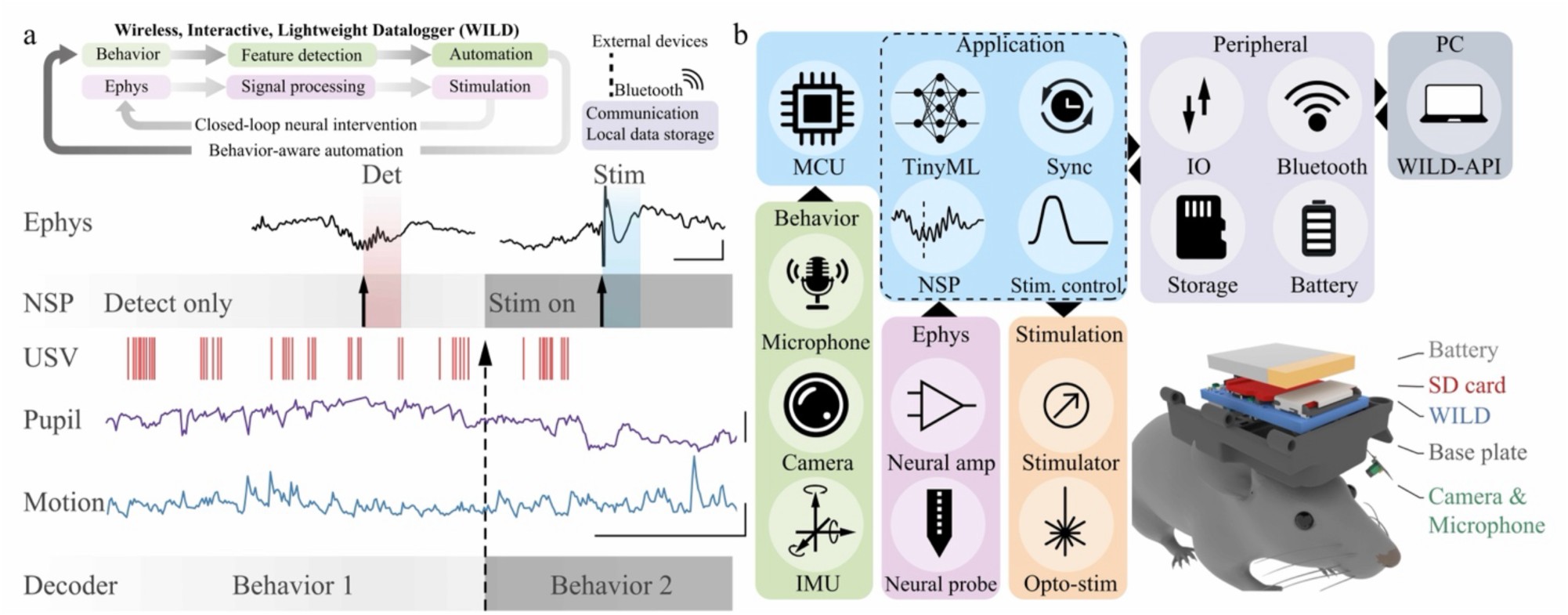
A wireless platform for recording and control of neural and behavioral activity in freely behaving small animals. **a)** Top: Schematic of WILD operations. Behavioral and electrophysiological signals are processed onboard to support feature detection, behavior-aware automation and closed-loop neural intervention. Data are stored locally, while wireless communication enables synchronization and monitoring through a remote device. Bottom: WILD enables wireless recording of neural electrical activity (‘ephys’; scale bar: 0.5 mV, 50 ms) alongside behavioral variables, including ultrasonic vocalizations (USVs), pupil dynamics, and locomotor parameters (scale bar: pupil diameter - ‘Pupil’, 0.2 a.u.; locomotion speed - ‘Motion’, 2 cm/s, 5 s), in freely behaving mice. **b**) System diagram of WILD. Behavioral and electrophysiological signals are fed into TinyML and neural signal processing (NSP) modules and stored in a microSD card. Real-time detection (‘Det’) of neural events and behavioral motifs is performed on board and can be used to control optogenetic manipulations in a closed-loop manner (‘Stim’). Communication with the host computer is achieved via Bluetooth, and seamless synchronization with external devices is provided, all controlled by WILD-API. The inset illustrates a mouse implanted with a WILD device, electrodes, battery, and protective 3D casing.

## Results

### A light-weight, multimodal, neural, and behavioral data-logger

The core element of our platform is a light-weight multi-modal data-logger (Fig. 1b, Extended Data Fig. 1a-b). While wireless neural recording systems have been previously developed^18,19^, our device overcomes existing barriers that have limited the broader adoption of these technologies and enables experimental paradigms that integrate multimodal sensing and on-board machine learning within an open-source framework.

Traditional wireless neural interfaces have usually been limited to the recording of neural activity, lacking other important functions, such as multi-modal sensing or on-demand manipulation. Our platform’s basic module features a 64-channel neural amplifier, a microSD card for data storage, and a Bluetooth module for wireless data communication up to 70 meters (Fig. 2a and Extended Data Fig. 1c). In addition, it includes additional sensors for monitoring behavioral variables. A 9-axis inertial movement unit (IMU) enables high-resolution motion tracking; a 320×320 pixels micro-camera captures pupil dynamics or animal point of view; and an ultrasonic microphone records animal vocalizations (Fig. 2a). All these different data streams are simultaneously recorded and seamlessly synchronized, together with external devices such as video cameras, through our open-source application programming interface (WILD-API), including an open-source graphical user interface (GUI) and command-line interface (CLI) (Extended Data Fig. 1d-e). We also provided documents for basic device operation command through Bluetooth low-energy traffic (Supplementary Table 2).

**Figure 2:**
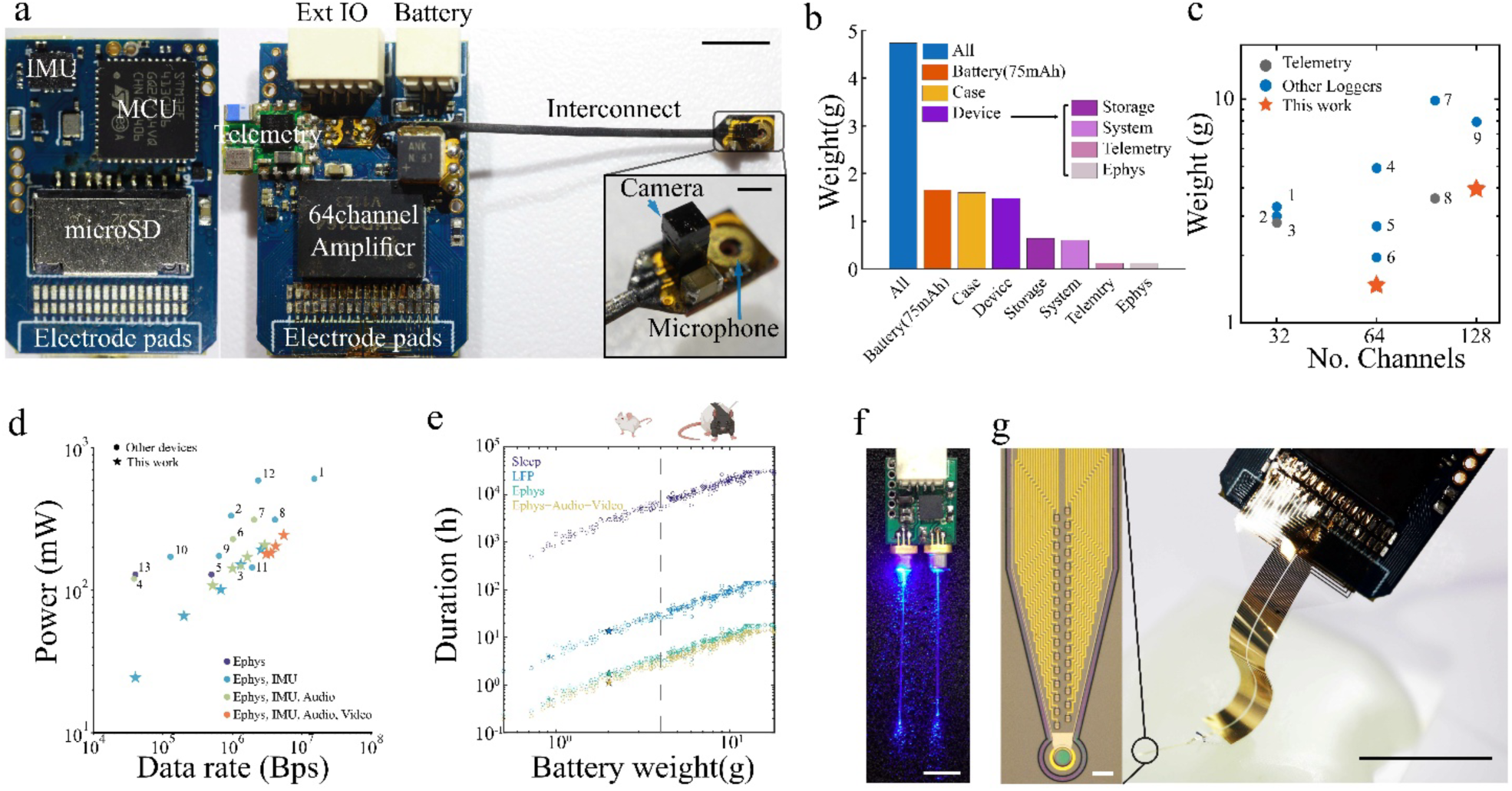
Device characterization. **a)** Picture of WILD basic module (scale bar, 5 mm). Inset: Zoomed-in picture of a multi-media capture module with embedded micro-camera and microphone (scale bar: 1 mm). IMU: Inertia measurement unit; MCU: microcontroller unit; Ext IO: extension Input/Output. **b)** Weight breakdown of the different components that integrate the device, including battery. Device itself weights < 1.5 g, reaching ∼4.5 g with case and battery. **c)** Weight versus channel count comparison of WILD (stars) with existing wireless neural interfaces^13,19–22^. See Supplementary Table 1 for detailed comparisons. **d)** Power efficiency (consumption versus data recording rate) comparison of different versions recording modes of WILD (stars) with existing wireless neural interfaces (dots, label correspond to reference number)^13,19–22^. **e)** Estimated operating times under power-saving idle mode for sleep, LFP (1250 Hz), only Ephys (20 kHz) and Ephys (20 kHz) with audio and video recording, using various lithium-ion batteries. Vertical dashed line indicates approximate weight limit for freely moving mice (∼ 5 g, including the battery. Note that the system provides ∼ 20 / 4 recording hours). Stars denote values measured with a 90 mAh battery. **f**) Implant-ready device with mounted flexible neural probe. Left: Microscopic image of high-density paralyne-C flexible neural probe (scale bar: 20 µm). Right: Photo showing mounted probe flattened on a phantom brain (scale bar: 10 mm). **g)** Picture of stimulation module with mounted laser diodes and optic fibers (scale bar: 5 mm). The weight of this module is 0.6 g.

Including these expanded functionalities, the total weight of the device is below 1.5 g, in a 23.3 × 15.7 mm circuit board (Fig. 2b). Compared to existing wireless neural recording and telemetry systems (that lack these advanced sensing and ML capabilities), WILD achieved the lowest weight (Fig. 2c), facilitating its use in freely-moving small animals such as mice or birds.

Another key element of WILD is a compact high-performance microcontroller unit (MCU, Extended Data Fig 1f). We developed an efficient signal acquisition pipeline that handles all data internally without the need for external memory or data processing modules, reducing device footprint and power consumption. As a result, WILD can record for longer periods of time than previous wireless recording systems (normalizing by channel count and battery capacity) (Fig. 2d,e, and Extended Data Fig. 1g-i). For example, a ∼2.5 g Li-polymer battery (130 mAh), that can be comfortably carried by a mouse, enables up to ∼3 hours of 64-channel recordings at 20 kHz (typical sampling rate for recording extracellular neuronal spikes) (Fig. 2e). For larger animals, such as rats, a battery of ∼5g would enable ∼9 hours recording (64-channels at 20kHz, Fig. 2e). In addition, the MCU incorporates embedded signal-processing and ML algorithms for real-time neural and behavioral event detection and generates trigger signals to control closed-loop optogenetics or electrical stimulation.

In addition to its embedded sensors, WILD is a modular, customizable platform that supports the expansion of channel counts (Extended Data Fig. 2a) or integration of additional modules through analog and digital interfaces (Extended Data Fig. 2b). An example of this is a miniaturized current stimulation module with integrated laser diodes and optic fibers for on-board optogenetic manipulations (Fig. 2f and Extended Data Fig. 2c). The combination of this stimulation module with WILD’s on-board real-time signal processing capabilities enables optogenetic stimulation triggered upon the detection of neural or behavioral patterns with sub-millisecond latency (Extended Data Fig. 2d-e), allowing precise real-time manipulations. All these functions can be flexibly configured through the WILD-API, enabling adaptable system operation for different experimental requirements (e.g., arbitrarily defined stimulation protocols, triggering with different neural or behavioral events).

Finally, WILD devices implanted in multiple animals can be synchronized through Bluetooth communications. Millisecond-precision synchronization was enabled by an onboard precision crystal oscillator and a two-way time-calibration process to ∼1 ms precision (Extended Data Fig. 2f-g). External systems could be synchronized through an expansion IO interface board (Extended Data Fig. 2h). This setup enables recording from multiple animals simultaneously, with all WILD devices and external input signals synchronized together.

WILD can interface with different common types of commercially available or custom electrodes through standard connectors (Extended Data Fig. 3a). To achieve maximal compactness and minimal weight with high-channel count electrodes, we microfabricated flexible neural probes based on a flexible parylene-C substrate and directly bonded them to the WILD board (Fig. 2f and Extended Data Fig. 3b). Each probe can be implanted independently, allowing targeting multiple brain regions simultaneously (Extended Data Fig. 3c). Moreover, these flexible probes eliminate mechanical mismatches between probe and tissue, enabling stable long-term single-unit recordings. Directly bonding them to the pads in the WILD circuit board eliminates the need for additional connectors, enabling probes, datalogger, and battery to fit inside a small 3D-printed casing (Extended Data Fig. 3d-f).

In summary, we have developed a fully integrated wireless, modular platform for energy-efficient monitoring of neural activity and behavioral variables, enabling real-time signal processing and closed-loop optogenetic manipulations in freely moving small animals such as mice.

### WILD enables multi-modal recordings during mouse social group behaviors

Mice are social animals that, in their natural habitats, live in groups with complex social dynamics^23,24^. However, the neural mechanisms of social behaviors in groups of freely interacting mice have been barely studied due to the limitations of current technologies. Tethered recordings severely restrict behaviors such as courtship or aggression and cannot be applied to multiple freely interacting animals due to cable entanglement (Supplementary Video 1). On the other hand, previous wireless neural interfaces have been limited in capturing behavioral signals together with high-bandwidth neural data, and have not been able to deliver responsive neural stimulation. To showcase how WILD enables experiments not feasible with previous approaches, we recorded groups of 2-4 male and female mice freely interacting in a large arena. Each mouse was implanted with a WILD module interfacing with a flexible neural probe and connected with a lithium-polymer battery (Fig. 3a).

**Figure 3:**
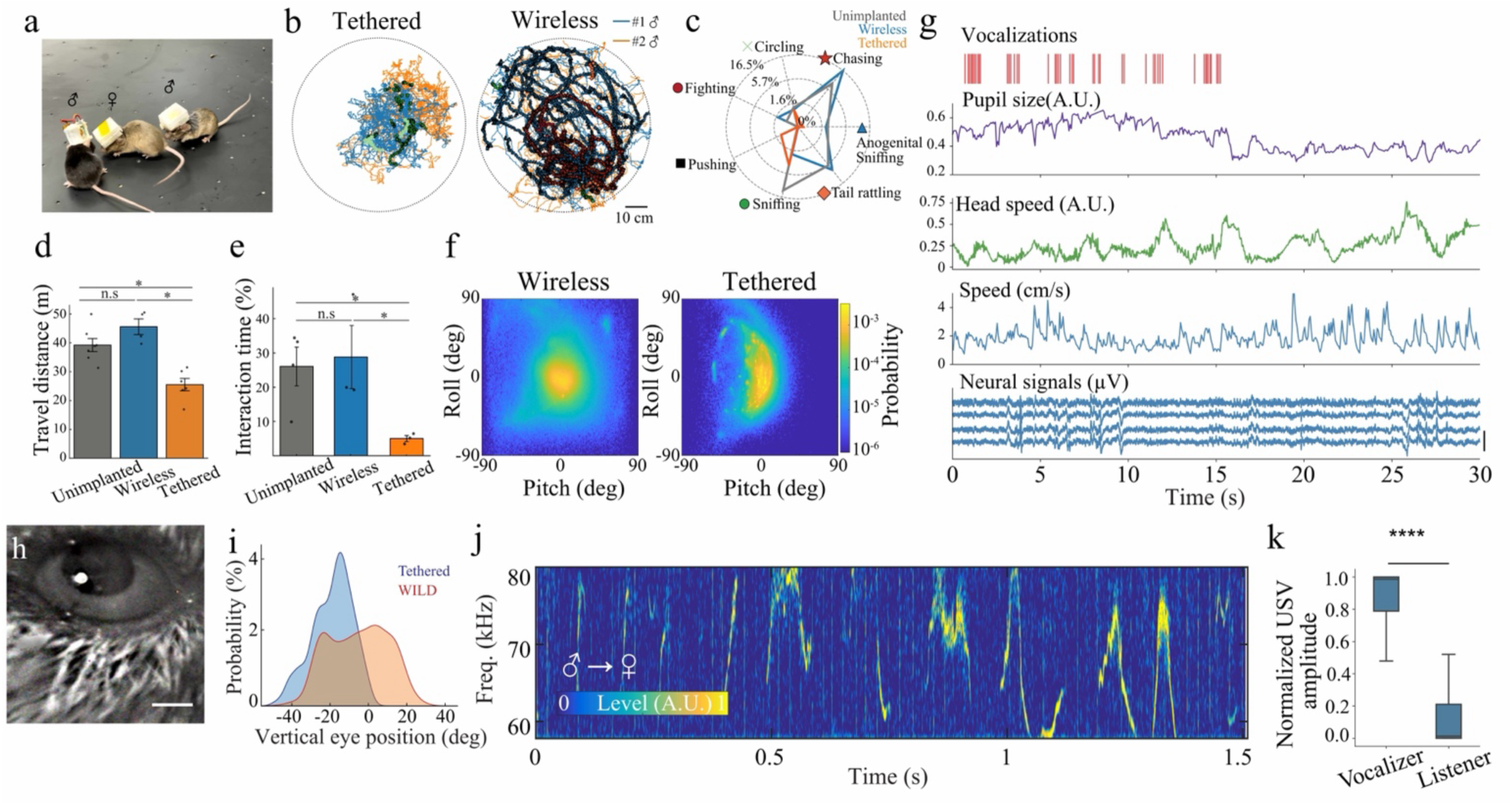
Multi-modal wireless recordings with WILD during mice’s unrestricted group social behavior. **a)** Picture of socializing mice implanted with WILD. **b**) Trajectories of two mice during free social interaction on an open maze during both tethered (left, using OpenEphys system) and wireless (right, using WILD) recording. Orange and blue traces represent the movements of two individual males. Different symbols mark the occurrence of different behaviors corresponding to Fig. 3c. Scale bar = 10 cm. **c)** Fraction of time of male mice with different recording conditions engaged in different social behaviors. **d**) Average travel distance of unimplanted (gray, n = 7 sessions from 2 mouse), wireless (blue, n = 4 sessions from 2 mouse), and tethered recorded mice (orange, n = 6 sessions from 2 mouse). Unimplanted vs wireless: p = 0.109, Mann-Whitney U test, p = 0.109, Neniamini-Hochberg False Discovery Rate(FDR); Unimplanted vs Tethered, p = 0.0023, Mann-Whitney U test, p = 0.007, FDR; Wireless vs Tethered, p = 0.0095, Mann-Whitney U test, p = 0.0143, FDR; Data are presented as mean values ± SEM. **e)** Male-male interaction time as a percentage of total session duration across different recording conditions Unimplanted vs wireless: p = 0.62, Mann-Whitney U test, p = 0.62, Neniamini-Hochberg False Discovery Rate (FDR); Unimplanted vs Tethered, p = 0.0114, Mann-Whitney U test, p = 0.0342, FDR; Wireless vs Tethered, p = 0.0249, Mann-Whitney U test, p = 0.0374, FDR, n = 10 sessions). Data are presented as mean values ± SEM. **f**) Comparison of head pitch and roll between tethered and wireless mice (roll/ pitch: wireless vs tethered p = 2.5527e-134 / p < 10^-100^, Mann-Whitney U test, n = 10 sessions). **g)** Example of multi-modal signals recorded from socially interacting mice. **h)** Mouse pupil captured by integrated micro-camera (Scale bar, 50 pixels). **i)** Distribution of vertical eye position during behavior (Kolmogorov-Smirnov test: D = 0.43, p = 1.54 × 10^-82^; Levene’s test for variance: F = 198.4, p = 5.97 × 10^-43^, n = 10). **j)** Example spectra of a male mouse USVs recorded with on-board microphone during social interaction with a female mouse. **k)** Relative amplitude for the same call as recorded by the microphone on the vocalizer and listener devices (n = 969 USVs, p < 5.769e-180, Wilcoxon test). Box-plots show median and 25^th^ and 75^th^ percentiles, with whiskers as the minimum and maximum values.

Compared with tethered conditions, mice implanted with WILD were more active, covered a larger area of the maze, and showed a larger spectrum of social behaviors, including sniffing, circling, chasing, and fighting (Fig. 3b-e, Supplementary Video 1). The percentage of time spent performing different social behaviors was similar between WILD-implanted and unimplanted mice, whereas tethered mice spent most of their time pushing away conspecifics (Fig. 3c,e). WILD-implanted mice traveled longer distances, displayed a larger variety of social behaviors, and prolonged interaction times compared to tethered mice, similar to unimplanted animals (Fig. 3b-e and Extended Data Fig. 4a-d). The on-board 9-axis IMU enabled 3D reconstruction of head movements for video-free behavior tracking (Extended Data Fig. 4e-g), which is particularly relevant during behaviors such as conspecific sniffing and fighting. WILD-implanted mice displayed a wider range of head pitch and roll than tethered mice, while mostly maintaining a central position not disputed by tether drag (Fig. 3f, Extended Data Fig. 4h). In addition, WILD-implanted mice showed significantly higher angular head mobility than tethered mice, suggesting WILD poses less head movement constraints (Extended Data Fig. 4i).

A key feature of WILD is that, in addition to neural recordings, it enables simultaneous acquisition of multiple behavioral variables (Fig. 3g). For example, mice emit different types of ultrasound vocalizations (USVs) during social interactions^25,26^. Pupil dynamics reveal changes in attention, arousal and brain state^27^, but they are difficult to monitor in the context of social interactions. WILD on-board microphone and camera allowed us to record USVs and pupil dynamics with high resolution during unrestricted social group behaviors (Fig. 3g). Our miniaturized head-mounted camera did not interfere with mice’s vision or eye movements, as evidenced by their wider range of eye movements compared to recordings performed with previously developed, tethered, head-mounted cameras^27–29^ (Fig. 3h-i and Extended Data Fig. 5a-e). WILD ultrasound microphone allowed for the detection of USVs during male-female interactions (Fig. 3j). Head-mounted microphones allowed us to differentiate between self-generated and conspecific USVs during unrestricted social interactions, something not feasible with over-head external microphones (Fig. 3 and Extended Data Fig. 5f-k). To verify this, we took advantage of the fact that during male-female interactions almost all USVs are produced by the male^30^. We found that most USVs showed high amplitude in the vocalizer (male) head-mounted microphone channel and near-zero amplitude in the listener (female) microphone channel (Fig. 3k). These simultaneous multi-modal recordings could thus enable dissection of the behavioral and neural basis of group social behaviors in mice.

To showcase the types of physiological insights that can be uncovered with this approach, we implanted mice with WILD devices and simultaneous flexible probes in the hippocampus and lateral septum (LS), important regions for social behaviors^31^. During unrestricted male-female interaction sessions in an arena, mice engaged in a range of different social behaviors (Fig. 4a). To visualize the heterogeneity of male behavior we extracted features of animal pose (derived from video) and WILD IMU, embedded them in a low-dimensional representation and color coded each interaction according to manually verified labels (Fig. 4b, left). Projecting USV rate, pupil diameter and hippocampal CA1 theta-band power (5-10 Hz) onto the same UMAP embedding revealed that these multimodal physiological and neural variables were differentially associated with distinct social behaviors (Fig. 4b). To explore this relationship in a more quantitative manner, we analyzed hippocampal CA1 - LS oscillatory theta coherence, pupil diameter and speed and USV rate around the onset of different behaviors (Fig. 4c and Extended Data Fig. 6). We found that ‘looking’ behavior was associated with increased CA1-LS theta coherence, larger pupil diameter, reduced pupil speed, suggesting relatively stable gazing, as well as a lower rate of USVs (Fig. 4c). During ‘approaching’ behavior, CA1-LS theta coherence increased, along with pupil diameter and USV rate. However, during ‘sniffing’, pupil diameter decreased whereas pupil speed increased accompanied by a higher USV rate. Finally, during ‘chasing’, we found a reduced CA1-LS theta coherence despite increased pupil diameter (Fig. 4c).

**Figure 4:**
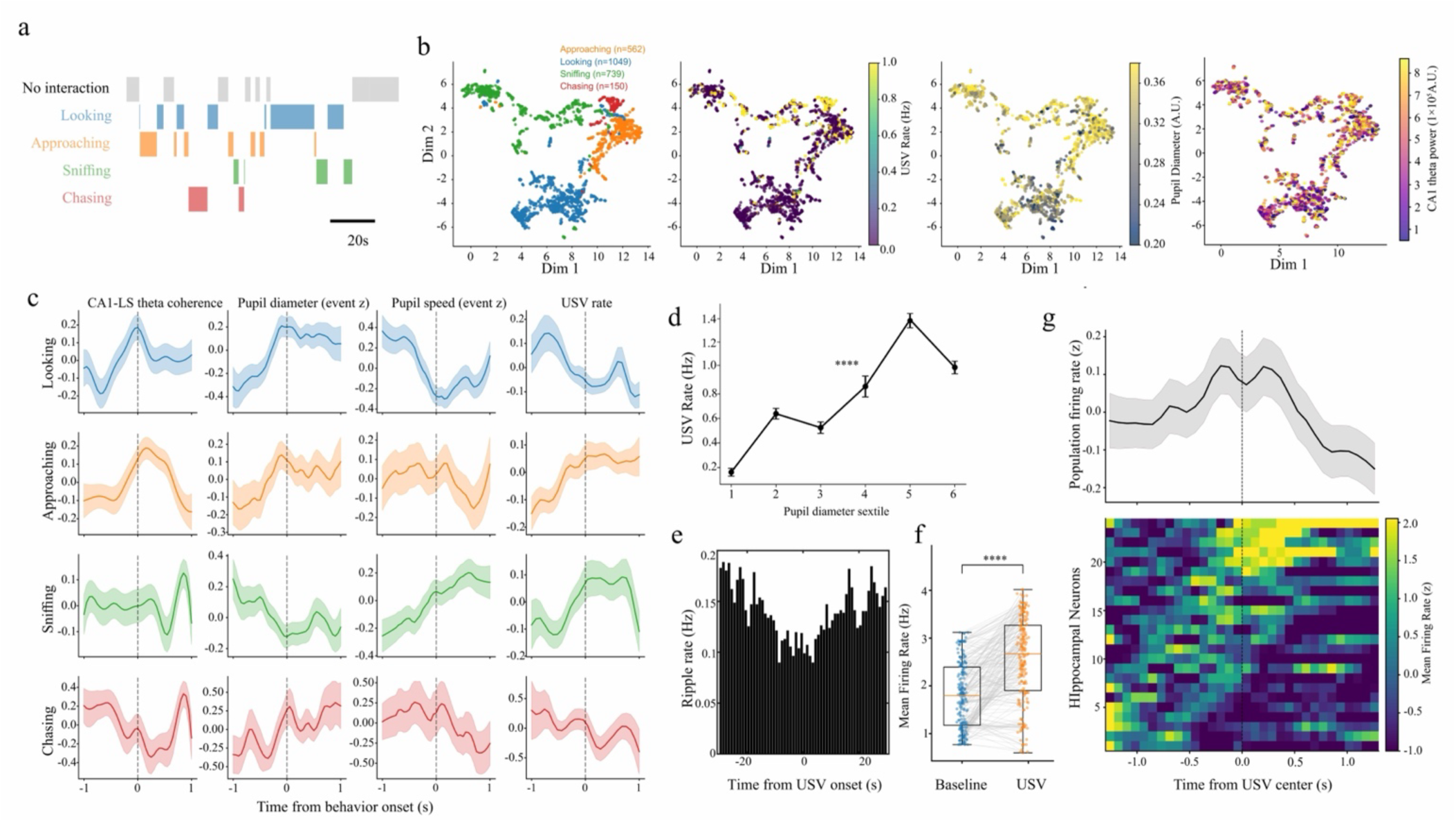
WILD reveals physiological correlates of different social behaviors. **a)** Ethogram showing distinct types of interactions during an example male-male interaction session. **b)** Left, 2D Uniform Manifold Approximation and Projection (UMAP) of body pose and IMU features in a representative male during male-female interactions. Color code shows manually annotated behaviors: approaching (orange, n = 562), looking (blue, n = 1049), sniffing (green, n = 739), and chasing (red, n = 150). Right, same embedding colored by USV rate, pupil diameter and hippocampal CA1 theta power. **c)** Peri-event modulation of neural and behavioral variables (columns) triggered by different social behaviors (rows) in males. Traces show the mean ± SEM from −1 to 1 s around behavior onset (0 s, dashed line) for looking (n = 70), approaching (n = 88), sniffing (n = 108), and chasing (n = 14) events (n = 2 mice). **d)** Mean ± SEM USV rate (Hz) is shown for each sextile of pupil diameter (r = 0.131, p = 1.47 × 10^-40^, n = 4,949 USVs from 6 sessions and 3 mice). **e)** Peri-event histogram of hippocampal ripple rate aligned to USV onset (stats and n). **f)** Average CA1 firing rate during USVs, restricted to low-speed non interaction periods (< 4 cm/s), and matched sample periods of non-USV (baseline) times (p = 3.73e-19, Wilcoxon signed-rank test, n=201 pairs, n = 24 neurons). Box-plots show median and 25^th^ and 75^th^ percentiles, with whiskers as the minimum and maximum values. **g)** Hippocampal CA1 neuron firing response around USVs (n = 201 events). Top, mean ± SEM population firing rate. Bottom, raster plot of individual neurons.

Overall, USV rate was strongly correlated with pupil size (Fig. 4d), suggesting that vocalizations are generated during transient states of increased arousal. We observed that most USVs coincided with periods of immobility or low-speed locomotion (Extended Data Fig. 6). During these periods, hippocampal activity is dominated by sharp-wave ripple (SWR) oscillations that strongly entrain neuronal activity. Surprisingly, USVs and SWRs were anticorrelated (Fig. 4e and Extended Data Fig. 6). Instead, hippocampal activity was selectively enhanced during USV events: CA1 neurons showed higher firing rates during USVs than during matched non-USV immobility periods (Fig. 4f). Furthermore, a subset of hippocampal neurons (∼40%) were strongly recruited during USV generation (Fig. 4g). Together, these results suggest that USV generation is associated with a distinct hippocampal activity state during immobility, distinct from SWRs, that selectively engages a subset of CA1 neurons.

### Neural and behavior recordings of groups of mice living in outdoor enclosures with WILD

Rodents in their natural habitats navigate and forage for food over long distances in complex environments. However, the neural mechanisms underlying these behaviors are typically studied in reduced laboratory mazes whose dimensions and complexity are limited by the use of tethers. To demonstrate the capabilities of the WILD system for the study of natural behaviors, we recorded rats and mice in very large outdoor enclosures (Fig. 5 and Extended Data Fig. 7). During fabrication, the WILD board was coated by a thin film of parylene-C, protecting it from humidity and mechanical damage, enabling animals to carry the device for multiple days while living outdoors, without compromising its function (Fig. 5a-e).

**Figure 5.**
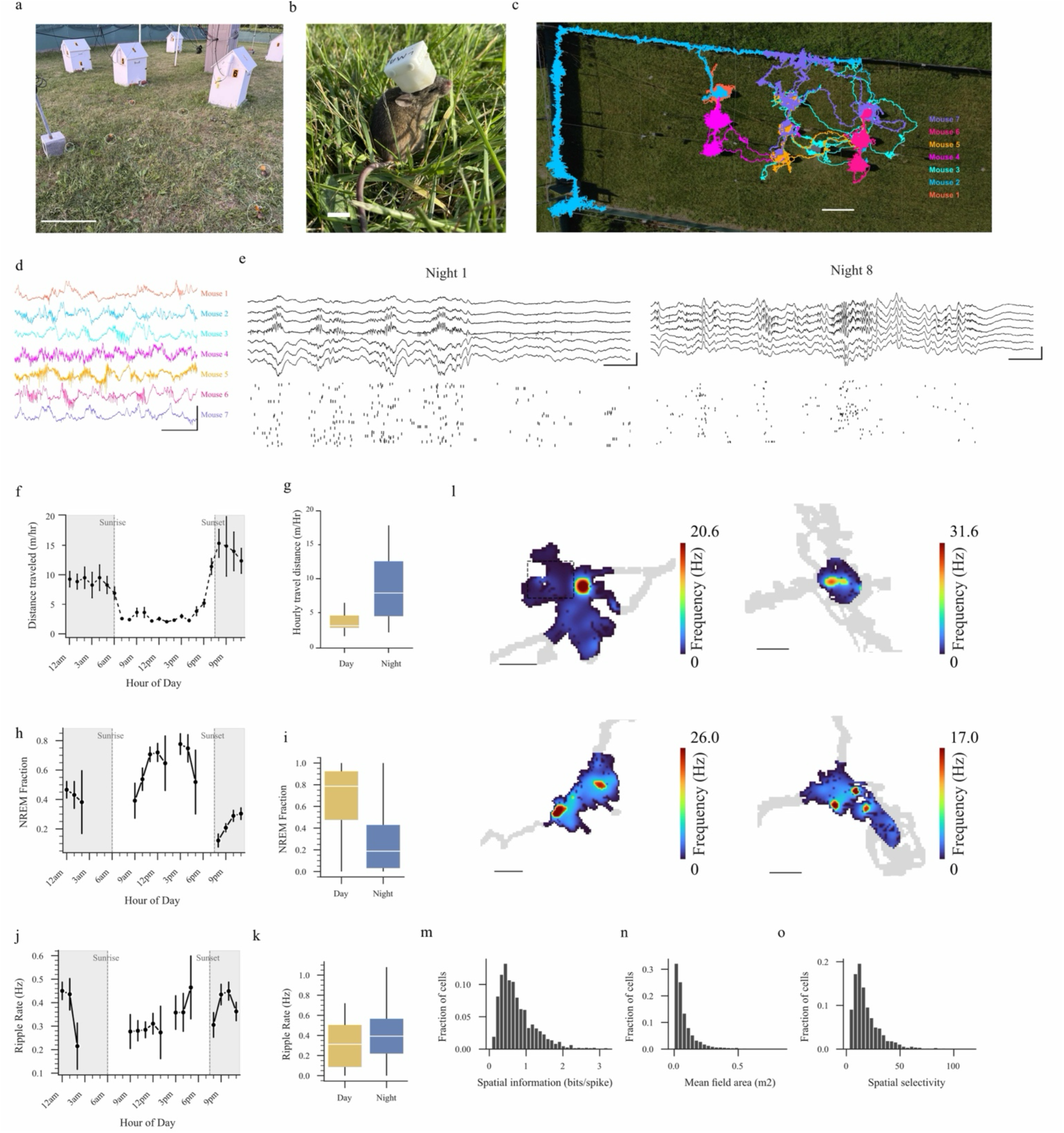
Wireless electrophysiology in freely moving mice in naturalistic outdoor settings. **a)** View of the semi-natural field enclosure with resource zones (white boxes). Six mice implanted with WILD devices are visible in the grass (white circles). Scale bar, 1m. **b)** Close-up of a mouse implanted with a silicon probe and WILD. Scale bar, 1cm. **c)** Trajectories of seven simultaneously recorded mice on night 6 from 7 PM to 7AM next day. Each color represents a different animal. Movements were recorded using WILD’s UWB module. **d)** Simultaneous hippocampal LFP traces from the same seven mice. Scale bar, 1mV, 200ms. e) Representative hippocampal LFP traces (top) and single neurons raster’s (bottom) from the same animal on night 1 (left) and night 8 (right), demonstrating recording stability of WILD. 1mV, 100ms. **f)** Mean distance travel per hour across a 24-hour cycle (n = 9 mice, nights 1-15). Gray shading indicates nighttime. Data are presented as mean values ± SEM. **g)** Distance traveled per hour was significantly greater during the night than during the day (day: 6.20 ± 0.71 m/h; night: 9.56 ± 0.91 m/h; n = 9 animals; p = 1.1 × 10⁻⁶, Wilcoxon rank-sum test). **h)** Fraction of time spent in NREM sleep across the day (n = 9 animals, 15 days). Data are presented as mean values ± SEM. **i)** Day vs. night NREM fraction. (n = 9 animals, p = 1.48 × 10^-17^, Wilcoxon rank-sum test). **j)** Mean SWRs rate during NREM sleep (Hz) across the day (n = 9 animals). **k)** Day vs. night NREM SWRs rate. (n = 9 animals, p = 0.0066, Wilcoxon rank-sum test). **l)** Example rate maps from four representative hippocampal place cells recorded in different animals in the enclosure. Examples were chosen to illustrate the heterogeneity of hippocampal place cells in the enclosures (e.g., with single or multiple place fields). Dashed boxes denote resource zones. Scale bar, 0.5m. **m)** Distribution of spatial information (bits/ spike) for all identified place cells (n = 1,636 cells from 9 animals). **n)** Distribution of mean place field area (m^2^) for all identified place cells (n = 1,636 cells from 9 animals). **o)** Distribution of spatial selectivity (peak in-field/ mean firing rate) for all identified place cells (n = 1,636 cells from 9 animals). Box-plots show median and 25^th^ and 75^th^ percentiles, with whiskers as the minimum and maximum values.

In one of these experiments, we implanted 9 male C57 mice with silicon probes in the dorsal hippocampus and WILD devices. Mice lived in and outdoor enclosure (38 × 15 m) for 15 days (Fig. 5a-b). Food was placed in 9 specific resource zones in the environment (wooden ‘shelters’; Fig. 5a). To investigate how mice explore a novel natural environment, characterize their spatial, social and circadian behaviors and their neural correlates, we tracked their behavior and recorded neural activity using WILD. Animal behavior was continuously recorded during this time using multiple cameras around the enclosure as well as an ultra-wide band position tracking module attached to the main WILD board (Fig. 5c). Neural recording quality and noise levels were not different from those obtained in the lab with standard tethered recordings (Fig. 5d and Extended Data Fig. 8), and remained stable while mice were living outdoors (Fig. 5e).

We first characterized mice movement patterns. There was a strong circadian modulation with mice becoming increasingly active shortly before sunset and remaining highly active until sunrise. (Fig. 5f-g). On average, mice travelled 6.20 ± 0.71 m/h during day and: 9.56 ± 0.9 m/h during the night (Fig. 5g). Conversely, mice spent a large fraction of the day sleeping, with long bouts of NREM sleep (Fig. 5 h-i). During NREM sleep hippocampal activity is dominated by sharp-wave ripples (SWR), synchronous network oscillations that support memory formation^32–34^. SWR rate displayed a surprising circadian modulation. It was relatively low during the day, despite the higher proportion of NREM sleep, and increased significantly during the night (Fig. 5j-k). This observation suggests an enhanced memory consolidation during brief periods of sleep during nighttime, perhaps due to increased amount of new information gathered during those periods.

During navigation, hippocampal ‘place cells’ fire at specific locations, encoding as a population an internal ‘map’ of the environment^35^. We were able to isolate and classify a large number of single neurons from these recordings (Fig. 5e and Extended Data Fig. 7) and identify multiple place cells. Place cells in the outdoor enclosure exhibit similar characteristics to those observed in much smaller laboratory mazes (Fig. 5l-o). Place field properties were heterogeneous across the population with some cells exhibiting multiple fields and others single ones (Fig. 5l), while the distribution of field size, spatial information and selectivity displayed a heavy-tailed distribution across the population (Fig. 5m-o).

Overall, this data demonstrates the utility of WILD to study the neural mechanisms of natural behaviors in rodents and other small animals (Fig. 5).

Despite its energy efficient design, WILD can only record at high bandwidth (64-channels, 20 kHz) continuously for ∼3-9 hours (depending on battery capacity, Fig. 2e), although recordings can be considerably longer at lower sampling rates (Extended Data Fig. 1 and 7). In some cases, it may be required to investigate behaviors that unfold over longer periods of time, during which retrieving animals to change batteries is not desirable. In order to support long-term, on-demand recording with minimal disturbance to the animal, we have designed the system with highly scalable power modes. When WILD is in idle mode, it is powered with a minimal power state (∼1mW), which could last for weeks with a 1.5 g battery (80 mAh, ∼300 hours). Switching between high-power (for 20 KHz, 64 channels recording) and low-power modes can be done by sending a remote command to the WILD module via Bluetooth or triggered by specific device events. This allows for restricting recordings to specific periods of active behavior, while saving battery life during prolonged periods of immobility and sleep.

### On-board real-time signal processing and TinyML enable neural and behavioral closed-loop manipulations

A powerful approach to investigate the neural mechanisms of behavior is to test the functional role of specific neural patterns or behavioral motifs by detecting them in real-time and delivering rapid, on-demand manipulations, such as optogenetic stimulation. This “closed-loop” approach offers a more precise control of neural circuit operations and enables linking specific neural patterns to behavioral outcomes^36,37^. Despite its advantages, its integration with wireless recording methods has been prevented by the challenge of real-time signal processing and stimulation delivery in embedded systems. Moreover, recent breakthroughs in the study of animal behavior have been catalyzed by the application of machine learning algorithms for the unsupervised classification of behavioral motifs, using either video^5,6^ or IMU^36,38^ data. However, these methods have not yet been deployed in embedded systems, preventing their integration with wireless platforms.

We tackled these challenges by developing a suite of real-time signal processing algorithms and efficient, light-weight ML models that can be executed directly by WILD on-board processor. Both neural and behavioral signals (IMU, sound, video) can be taken as inputs, enabling the rapid detection of transient neural activity patterns, specific behavioral motifs, or a combination of both. We have implemented three parallel on-board signal processing pipelines for detecting neural and behavioral events in real-time. The first pipeline implements low-latency digital signal processing (Fig. 6a). This allows performing different operations on streamed neural signals, such as filtering in specific frequency bands and estimating the phase of an oscillation or calculating the time-varying power of a signal, such as neuronal spiking. Logical operations, such as power threshold crossing or detection of a specific oscillation phase, can be implemented and used to enable the triggering of a stimulation module. These operations can be performed on multiple neural signals simultaneously, for example, to detect synchronous events across different brain regions. Our custom algorithms enable high-accuracy signal detection with a latency of ∼5 µs (Extended Data Fig. 9a-h), achieved by eliminating additional processing delays from data communications.

**Figure 6:**
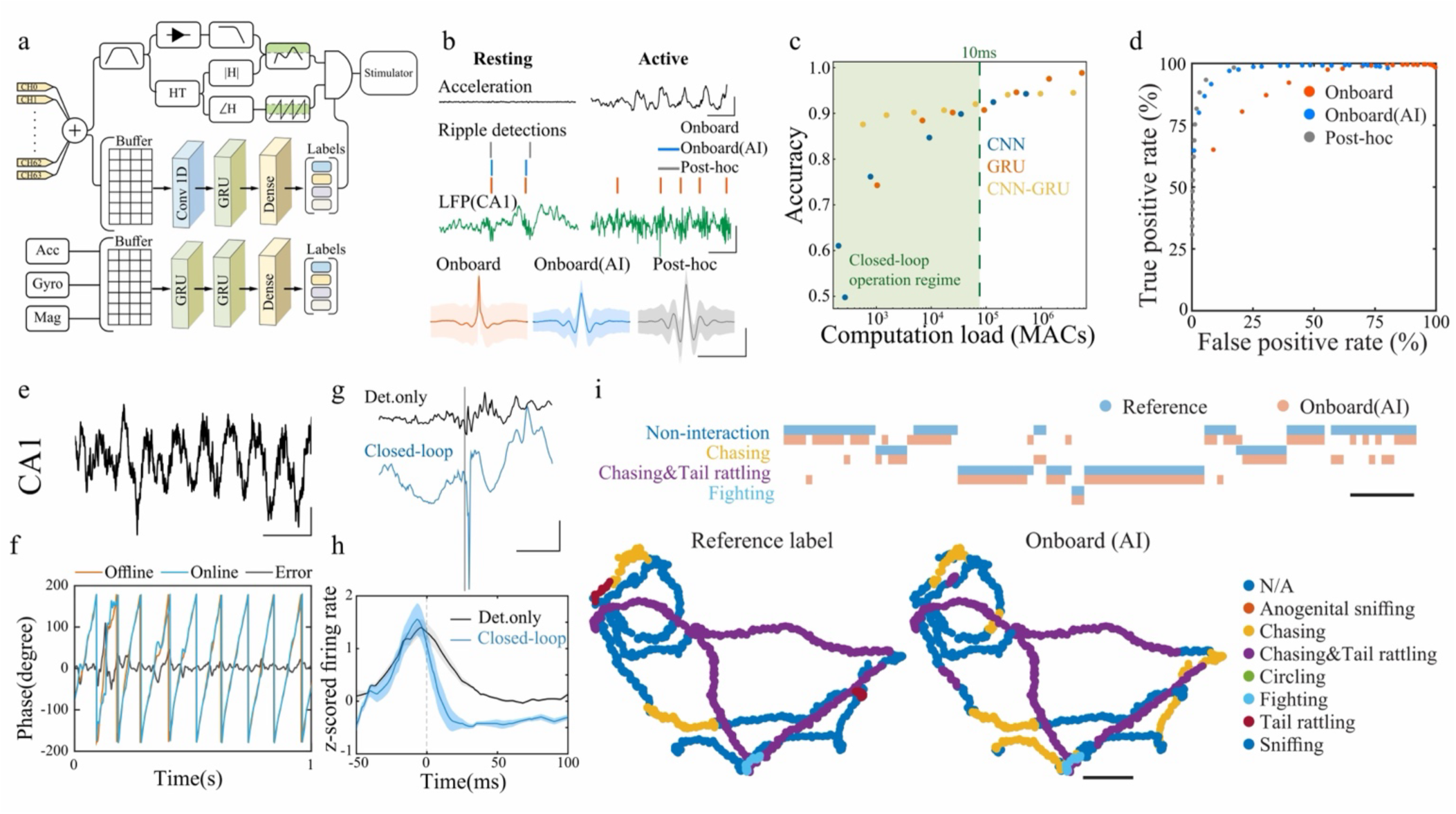
Neural and behavioral closed-loop manipulations with WILD in behaving mice. **a)** Schematics of embedded real-time signal processing. Neural signals are processed by a low-latency digital signal processing pipeline (top). Neural signals are also cached and analyzed by an embedded TinyML model (bottom). Outputs from these pipelines are used to control stimulation. **b)** Example of SWR detection in a mouse during sleep (left) and behavior (right). Vertical lines indicate real-time SWR detection with only on-board signal-processing (orange), with added TinyML denoising (blue), and post-hoc (gray). Green trace is hippocampal LFP. Note spurious SWR detection during behavior due to EMG noise (right). Bottom panels are average traces of detected events. **c)** Estimated accuracy of TinyML noise identification model compared with model size and structures. Target inference times are indicated by a dashed line. **d)** ROC curve of SWR detection without (orange) and with on-board denoising (blue), alongside post hoc ripple detection (gray). (nSPW-R=2,378). Paired bootstrap: Online AUROC 0.717 vs AI AUROC 0.844; Delta AUROC −0.127 (95% CI −0.162 to −0.092), p = 5.96e-12 paired bootstrap Δ area under ROC; Online − AI. **e)** Example trace of CA1 LFP during theta oscillations, **f)** Example of onboard theta phase estimation compared with an offline phase based on Hilbert transform. **g)** Example of optogenetic ripple closed-loop interruption. Black trace shows a detection-only trial; blue trace shows a closed-loop optogenetic interruption trial. Gray line indicates the time of detection (Scale bar: 50 ms, 0.5 mV). **h)** Firing rates of CA1 units (n = 7) during SWRs in detection-only (n = 20,472) and closed-loop stimulation (n = 10,085) conditions. Data are presented as mean values ± SEM. **i)** WILD real-time behavior prediction. Upper panel: ethogram of online prediction (orange) compared with manually labeled behavior (blue). Bottom panel: animal trajectory during a male-male chasing event, colored by online-predicted behavior labels.

The other two signal processing pipelines are TinyML engines for real-time inference from pre-trained artificial neural network models for neural pattern and behavioral motif classification, respectively. These TinyML models are designed to be computationally efficient, minimizing inference time and power consumption while achieving desired accuracy. The accuracy of different neural network architectures scales differently with model size and is highly dependent on the type of input data. We found an optimal tradeoff between accuracy and latency for neural signal inference by combining a convolutional neural network (CNN), for sub-sampling, and a gated recurrent unit (GRU), to capture signal temporal features (Fig 6b, Extended Data Fig. 9b). Running this model on WILD achieved a > 90% accuracy in neural pattern detection with a latency below 10 ms (Fig. 6c).

We performed several experiments to demonstrate on-board signal processing and closed-loop stimulation capabilities of WILD. Sharp-wave ripples (SWRs) are transient (50-100 ms), high-frequency (∼150 Hz) oscillations that occur during pauses in behavior and sleep. They have been widely studied for their important roles in learning and memory^32,39^. Fast and accurate neural interventions are required to investigate the causal relationship between SWR occurrence and behavioral outcomes^31,34,40,41^. This is notoriously difficult in behaving animals due to the confounds introduced by other sources of high-frequency activity, such as electromyogram (EMG) activity, that contaminate hippocampal neural recordings^42,43^. We performed online SWR detection on a mouse with large EMG contamination (Fig. 6d). WILD TinyML model was trained to reject EMG activity, which was achieved by introducing minimal processing delay (Fig. 6b,c). After denoising, real-time SWR detection improved significantly, reaching a performance similar to post-hoc detection (which uses multiple non-causal and computationally demanding processing steps) (Fig. 6d).

To demonstrate that WILD can enable experiments that are not feasible with current methods, we performed neural closed-loop manipulations in mice while conducting a spatial foraging task in a 3D maze. Mice started the trial above an elevated platform and had to find their way down and into an entrance to the structure underneath, where a food reward was hidden (Extended Data Fig. 10c). Transgenic mice expressing channelrhodopsin (ChR2) in pyramidal cells were implanted with silicon probes and optic fibers in the hippocampus and prefrontal cortex, and a WILD device with an optogenetic stimulation module. During this task, two main network activity patterns were present: SWRs, which dominated during periods of immobility, and a slower, sustained “theta” frequency (∼8 Hz) oscillation during locomotion. We performed two types of optogenetic manipulation experiments. First, we detected periods of sustained theta oscillations and computed their real-time phase, then delivered optogenetic stimulation restricted to a specific phase (Fig. 6e-f). In control conditions, pyramidal cells were strongly phase-locked to theta oscillations, but this phase-locking was disrupted by our manipulation. We also detected SWRs and disrupted them by delivering short blue-light pulses (Fig. 6g), which abolished SWRs and silenced the associated neuronal firing (Fig. 6h). This experiment demonstrates how WILD enables rapid and selective manipulation of specific neural patterns.

In another experiment, we performed real-time behavioral pattern classification in pairs of freely interacting male mice. The goal was to identify specific behavioral motifs in real-time in conditions where video classification would be challenging (e.g., groups of interacting mice in cluttered environments) to enable closed-loop manipulations, all in a fully wireless manner. First, we labelled different social behaviors of previously recorded videos. Then we used these behavioral labels to train WILD’s on-board TinyML model. The TinyML engine implements a behavior classifier, based on IMU signals, that can detect specific, transient behavioral motifs or “syllables” (Fig. 6a). We obtained the best performance with a deeper GRU architecture (Extended Data Fig. 10). Finally, we used the trained model to detect different behaviors in real-time. The low latency of our on-board behavioral prediction (>100 ms) allows using it to trigger optogenetic or other types of manipulations to disrupt neural activity during specific, transient behaviors, even in groups of mice during unrestricted interactions. Performance of real-time behavioral identification was highly similar to the ethogram from manually labelled video (Fig. 6i), demonstrating the accuracy of WILD on-board behavioral syllable prediction and its suitability for closed-loop manipulations.

## Discussion

We developed a modular wireless, responsive neural interface platform (WILD) that overcomes key limitations of existing technologies, enabling experiments that are not feasible with current methods for investigating the neural basis of natural behaviors in small animals. To facilitate its adoption by the community, we have made all the hardware and software elements of this platform open-source and fully documented online.

Recent advances in tethered neural interfaces have enabled large-scale neural recordings in mice and other small animals^2,18,44^. However, this approach is not compatible with a wide range of naturalistic behaviors, especially those involving multiple animals, large-scale spaces, or cluttered environments. These limitations can be overcame using wireless devices^14,15,45,46^. However, existing wireless interfaces lack important capabilities such as on-board real-time signal processing or simultaneous multimodal sensing, and have limited recording bandwidth and duration under weight constraints as those imposed by freely moving small animals.

The WILD platform offers several other key advantages over previously published and commercially available systems. First, accessibility: WILD is constructed entirely from off-the-shelf components and is compatible with standard neural probes, including flexible probes, silicon probes, and tetrodes. WILD uses Bluetooth Low Energy for wireless communication, allowing operations without additional expensive equipment while still supporting communication over tens of meters and millisecond-level synchronization. For applications requiring longer range, WILD can be readily adapted to alternative wireless protocols, such as LoRa, sub-GHz radio, or cellular networks. Second, functional integration: whereas existing wireless neural interfaces typically support only one recording modality (Supplementary 1), WILD enables concurrent or selective acquisition from a diverse set of sensors, with reserved connectivity for future expansion. Third, real-time event detection: WILD incorporates a standalone signal-processing pipeline that operates on both electrophysiological and behavioral inputs, enabling detection of neural events, as well as model-based signatures derived from neural and behavioral signals. Unlike existing systems, WILD supports concurrent closed-loop optogenetic (or electrical) stimulation and high-fidelity signal acquisition within a fully wireless architecture. These design features further enable high-performance real-time neural signal processing and closed-loop manipulation in untethered animals, including in environments beyond conventional laboratory settings. At the same time, WILD remains more lightweight and more power-efficient than other existing wireless recording devices due to its integrated system architecture and optimized data handling. The possibility of switching between high-power and idle recording modes on demand (e.g., triggered by animal movement) enables prolonged recordings without the need for battery change. Wireless charging via photovoltaic or electromagnetic coupling^47,48^ could also be used to extend operating time.

Compared to existing technology, our platform offers advantages in high-performance real-time neural signal processing, enabling multi-modal recordings and closed-loop neural manipulations in groups of untethered animals, even outside conventional laboratory environments. In contrast to tethered or head-fixed approaches, our platform enables circuit-level investigations of group interactions, courtship, aggression, and navigation in large environments in small animals. To illustrate some potential applications, we demonstrated how mice carrying WILD devices behave in a similar way to unimplanted animals during social group behaviors, and how simultaneous recording of neural and behavioral signals uncovers previously unobserved correlates of social behaviors. Our recordings of mice living in an outdoor enclosure also allowed to observe rodent place cells in a natural environment. This approach can thus be extended to many other types of experiments of social, spatial and other types of behaviors in a variety of animals, from rodents to birds.

Onboard ML models support real-time detection of behavioral and neural signatures. We illustrated this by performing optogenetic manipulations of specific neural patterns in mice behaving in a 3D maze, and by the real-time detection of specific behavioral motifs during social interaction. While these types of experiments have been performed before, they required tethered setups and external computers for signal processing^36,34,27,49^. Implementing this capability in a wireless system and integrating it with onboard optogenetic stimulation enables applying it to a much wider range of behavioral paradigms that are incompatible with tethered approaches. Furthermore, the development of embedded TinyML on our platform aligns with growing industrial interest in edge computing^50^. We anticipate that continued hardware advances will bring even greater computational capacity to compact processors that, deployed in neural interfaces such as WILD, can further expand their capabilities for investigating the neural mechanisms of natural behaviors both inside and outside the laboratory.

## Acknowledgments

The authors thank members of the Oliva and Fernandez-Ruiz labs for providing useful feedback on the manuscript and Michael Sheehan for logistic assistance with outdoor experiments.

## Author contributions

Z.Z. and J.P. fabricated the devices, Z.Z. developed the software, Z.Z., H.C., C.L., P.P., M. A., A.O. and A. F-R. performed the experiments, Z.Z, H.C. and P.P. analyzed the data, A.F.R, Z.Z. and A.O. wrote the manuscript with input from all authors. A.F.R. and A.O. supervised the work.

## Methods

### Device fabrication

Circuit design is performed in an EDA software (EaglePCB, Autodesk). PCBs are fabricated with an online vendor (NextPCB). The device is assembled manually with a reflow station and a soldering station. Assembled devices are programmed through USB device firmware update mode and tested for power consumption and function integrity. We used a high precision quartz oscillator for the clock (ABM11, Abracon; ±10ppm), then the crystal load capacitor is manually trimmed to achieve <1ppm clock accuracy (tested with Time & Frequency Analyzer, Moku:Go, 0.78ps resolution). Connectors are first protected with Kapton tape or a dummy plug, then a hermetic protection layer is formed with 3μm parylene-C deposition. The device is then prepared differently for standard neural connectors (A79025, Omnetics Connector Corp) or directly bonded with a flexible neural probe. We designed the flexible bonding pattern to be the same as the neural connector landing pattern. A neural connector can be soldered to the electrode pads to be compatible with commercial silicon probes; a thin layer of low-temperature solder (SMDLTLFP, Chip Quik) is first prepared on the electrode pad with a 0.1mm stencil. The flexible neural probe was aligned to the electrode pads using a wet brush, then pre-baked at 80 °C in a constant-temperature oven for 5 minutes to evaporate residual water. The temperature was then raised to 160 °C for 10 minutes to melt the solder and form the electrical connection. The quality of the probe is checked by electrode impedance measurement in saline solution, with Ag/AgCl electrode as a reference. Laser diode (PL450B, ams-Osram) of the opto-stimulation board is coupled to the 200μm optical fiber with two micro-manipulators help adjust the angle and offsets, and fixed with UV-curing optical adhesives (Optical Adhesives, Norland) when the maximal optical output position is found.

### Flexible neural probe fabrication

Flexible neural probes are fabricated in standard cleanroom facilities. Photolithography masks are designed with EDA software (L-edit, TannerEDA). Lithography of 5’ Cr masks is performed with a laser writer (DWL2000, Heidelberg Instruments), followed by developing and etching (HMP900, Hamatech). 4’ silicon wafers are first cleaned with a mixture of hot Sulfuric Acid and Hydrogen Peroxide (Hot Piranha, Hamatech), then thoroughly washed with IPA, acetone, and DI water spin rinse. A 1.5μm layer of parylene-C was then deposited with PDS 2010 LABCOTER. The thickness of deposited parylene-C was measured by a stylus profilometer (DektakXT, Bruker). We used a lift-off process for metal patterning. Negative photoresist (AZ nlof 2020, MicroChemicals GmbH) is spun at 3000 RPM, 30s, followed by 110°C soft-baking for 60s. Photolithography is performed with a contact aligner (MA6-BA6, SUSS MicroTec), followed by 110°C post-exposure baking for 60s. Photoresist was then developed with TMAH-based developer (AZ726MIF, MicroChemicals) for 90 seconds. 20nm Ti followed by 150nm Au is then deposited with an E-beam evaporator (Mark 50 E-beam Evaporator, CHA industries). Metal lift-off is performed by dipping the sample in MICROPOSIT™ REMOVER 1165 for 24 hrs. Following metal lift-off, the sample was thoroughly cleaned before spin-coating a negative photoresist (AZ nLOF 2020, MicroChemicals GmbH) to define the Pt electrode pattern on top of the Au interconnects. A 50 nm Pt layer was then deposited using the same E-beam evaporator (Mark 50 E-beam Evaporator, CHA Industries), followed by lift-off under the same conditions as used for the Au metal pattern. Optical inspection was performed to verify the integrity of both the Au interconnects and the Pt electrode pattern. A second parylene-C layer is deposited to form full encapsulation. A thick photoresist mask (MEGAPOSIT SPR220, DuPont) is then spin-coated (1500rpm, 30s) onto the wafer. Wafer is first soft-baked (90°C,30s; 115°C, 6mins), rested at room temperature for 20mins, then exposed with a contact aligner. Wafers then rest for >1Hr, followed by post-exposure bake (90°C, 30s; 115°C 3mins). Wafers were then put on a wafer holder to cool to room temperature, followed by development (AZ726MIF, 90s). We used oxygen plasma etching to perform anisotropic etching of the electrode openings and probe outlines (Minilock III ICP etcher, Trion Technology). A mixture of high-conductivity PEDOT:PSS solution was prepared by mixing 10 g of PEDOT aqueous dispersion with 4 g of sorbitol. The mixture was thoroughly mixed and then sonicated for 30 min to improve homogeneity. After sonication, the solution was filtered to remove particulates. Immediately before use, 60 µL of 1% GOPS solution was added as a crosslinker, followed by one drop of DBSA. The final solution was mixed gently before coating^51^. PEDOT:PSS is two-step spin-coated (500 rpm 10s/1000 rpm 60sec) on the wafer, followed by baking (115°C, 20 mins). A protective layer of photoresist is patterned for PEDOT:PSS patterning (AZ nlof 2020) as mentioned above, followed by oxygen plasma etching. Surface conductivity of the wafer is monitored to evaluate if the excess PEDOT:PSS is removed. To remove the photoresist used for the PEDOT patterning, each sample was individually immersed in AZ NMP Rinse and subjected to ultrasonic agitation at room temperature for approximately 5 seconds to ensure complete removal of the PR. Successful photoresist removal was typically indicated by a color change of the NMP Rinse to yellow. The sample was then rinsed with DI water. While submerged in DI water, the surrounding Parylene-C film deposited on the wafer was carefully peeled off using fine tweezers. Subsequently, localized water flow was gently applied using a pipette along the edge of the probe to initiate delamination from the wafer. Upon release, the probe floated to the water surface. The floating probe was then carefully guided and aligned to the PCB Pad using a fine brush. Due to its thin and delicate structure, tweezers often caused tearing, so alignment was performed entirely with the brush. After alignment, the assembly was dried on a hot plate at 80 °C for approximately 10 minutes and then soldered to a PCB board using a low-temperature solder paste (melting point: 138 °C).

### Wireless Recording with WILD and synchronization with external devices

During recording, WILD devices maintain an active Bluetooth connection with the host PC, enabling full bidirectional communication. PCs equipped with Bluetooth Low Energy (BLE) modules were used to control the WILD devices. The host PC clock relies on an internal crystal oscillator, which is subject to drift over time. To minimize this, we calibrated the PC clock against an online reference using an open-access tool (Time Calibrator, Fountain Computer). A custom firmware was implemented on the WILD devices to report their internal clock values to the PC via serial communication, allowing us to compare device and host timestamps and quantify timing differences (Extended Data Fig. 2). Upon connection to the PC, each device underwent a 20-second clock calibration procedure to synchronize its internal clock with the host PC clock. After synchronization, the PC controlled the device through a set of commands, including initiating and stopping recordings, previewing signals, adjusting device parameters, managing external I/Os, and performing impedance measurements. The source code for the WILD PC API is provided (https://github.com/ayalab1/Neurologger), allowing users to integrate the system with external behavioral control or tracking software. Communications from the WILD device are processed through event-driven handlers, which can be linked to the APIs of other experimental control systems. Additionally, an I/O expansion board (Extended Data Fig. 2j) enables integration with external experimental hardware by providing digital input and output lines for synchronization and control. This interface allows external devices to trigger WILD operations or receive selected internal signals in real time, facilitating integration with existing behavioral control and tracking systems.

High-speed Bluetooth communication allows a single host computer to connect to up to seven WILD devices simultaneously (see Fig. 5). This configuration enables real-time communication with all connected devices, including signal streaming, transmission of control commands, and other online operations. Alternatively, a larger number of devices, up to a few tens to one hundred devices, can be managed by the same host computer through sequential connections. In this mode, continuous real-time communication with all devices is not maintained, which limits certain online operations. However, during the initial connection, each device’s internal clock is synchronized with the central PC clock. This synchronization ensures that recordings remain temporally aligned across devices throughout the entire recording session, even when continuous real-time communication with the host computer is not maintained.

We developed a continuous two-way BLE delay and clock-offset calibration process to align the WILD device real-time clock (RTC) with the host PC time base. During each BLE timestamp exchange, the PC recorded the packet transmission time and response arrival time, while the device recorded packet reception and response time using its onboard RTC. These timestamps were used in a Network Time Protocol-style calculation to estimate the PC-device clock offset and BLE round-trip delay. The calibration was repeated before and during recording to reduce the effect of BLE scheduling jitter and to correct residual RTC offset and drift.

Application-level BLE communication latency was estimated after clock-offset correction. The PC sent a timestamped BLE packet, the device recorded the packet reception time using its RTC and returned this timestamp to the host. The host then calculated the communication delay from the corrected difference between PC packet transmission time and device packet reception time. This measurement reflects effective real-time BLE latency at the application level, including PC scheduling, BLE stack timing, connection interval effects, firmware processing and timestamping overhead, rather than physical-layer radio propagation delay.

External behavioral events were detected using a GPIO-USB interface board and timestamped directly on the host PC using the PC system time base, rather than through BLE. Because the WILD RTC was calibrated to the PC time base, PC-recorded GPIO event timestamps could be aligned with device-recorded neural, behavioral and system timestamps.

Different operations can be preprogrammed before the start of recording or controlled online through Bluetooth commands. In addition, onboard signals such as those from the IMU can be used as real-time triggers for device actions. Typical operations include remote control of recording start and stop, as well as conditional operations based on behavioral signals. For example, recording can be initiated or the device can enter sleep mode based on animal movement detected from online IMU activity. A summary table of commonly supported command operations is included in Supplementary Table 2.

WILD supports multiple communication interfaces for device configuration, electrode placement, and recording. The system is equipped with general-purpose I/O (GPIO) lines that can be used for synchronization with external devices, control of stimulation circuits, or communication with other experimental hardware. For high-bandwidth data transmission, WILD includes a USB 2.0 interface that enables real-time streaming of signals to a host computer. This interface is typically used during procedures such as electrode implantation, when high-resolution electrophysiological signals need to be monitored to guide electrode placement. To facilitate camera positioning during surgery, we provide a custom Python script that enables live streaming of the head-mounted camera images. In addition, local field potential signals from selected channels can be streamed to the host PC through the Bluetooth interface simultaneously, allowing wireless monitoring during recording sessions.

### Experimental model

All experiments conformed to guidelines established by the National Institutes of Health and have been approved by Cornell University’s Institutional Animal Care and Use Committee. All rats/mice were kept in the vivarium on a 12-hour light / dark cycle with *ad libitum* access to food and water except during training. Temperature and humidity in the room were kept at 68-72 °F and 40-60%, respectively. Animals were housed with a maximum of 2 per cage before surgery and changed to individual housing afterward. 11 mice were used for electrophysiological recordings. Long-Evans rats (male and female, 3-8 months old) were used for a 128-channel electrophysiology experiment. Male B6FVBF1/J mice (3-6 months old, 27-32 g; The Jackson Laboratory) were used to compare tethered, unimplanted, and wireless conditions. CaMKIIα-Cre/+; Ai32/+ mice (approximately 25-30 g, 3-6 months of age) were used for optogenetics experiments.

### Surgical procedures

Electrophysiology implantations used flexible neural probes for mice and silicon probes for rats; For flexible neural probe implantation, mice were anesthetized with isoflurane. Skin on top of the skull is disinfected, opened with a scalpel, and tissue on the skull is removed, then the skull is cleaned with hydrogen peroxide. Two stainless ground screw are tapped on the rear side of the lambda. A thin layer of strong dental cement (C&B Metabond, Parkell) was applied on the surface of the skull, excluding the planned craniotomies windows. A 3D printed base plate is fixed to the skull with Metabond. Craniotomies were performed based on stereotaxic coordinates of hippocampus CA1 (−2.0 mm AP, +2.0 mm ML). After dura removal, flexible probes are flattened on the brain surface. A chemically sharpened 50μm tungsten wire (California Fine Wire) was used as an insertion tool to guide the probe for penetrating brain tissue, with a slow insertion speed (1mm/min). After reaching target depth, the tungsten wire is retracted, leaving the flexible probe in place. For the implantation with optical fiber, the optical fiber is inserted, followed by probe insertion with the same coordinates. Craniotomy windows are first protected with artificial dura-gel, then covered with a 1:1 mixture of low-temperature paraffin and paraffin oil. Micro camera is positioned towards the pupil and validated with live imaging. The device is then fixed to the base plate with dental cement. Post-operative care included administration of carprofen subcutaneously for 3 days, and the skin near the implant was treated with triple antibiotic ointment. For rat implant, two high-density silicon probes (Diagnostic biochip) mounted to a microdrive are used for each animal, with coordinates for bilateral hippocampus CA1 (−4.0 mm AP, ±3.0 mm ML). Animals were placed in their home cage over a heating pad for post-surgical observation and were allowed to fully recover for>7 days post-surgery. For silicon probe implantation, electrode depths were gradually adjusted until the hippocampal layers were identified with electrophysiological signatures.

### Social interaction experiment

Trials were conducted in a 31-inch, disk-shaped open-field arena. The ambient lighting in the room was adjusted to ensure uniform illumination. An overhead camera (aca1300-200uc, Basler AG; 40 fps or acA720-520uc; 24.9 fps) was used to record videos of animal behavior. Devices were synchronized over Bluetooth prior to the session start. Video and multi-modal recordings were initiated separately, with start times logged for alignment. To begin each trial, an implanted mouse was placed in the open-field arena for 5 minutes to familiarize itself with the environment. A second animal was then introduced to the arena and allowed to interact socially with the first animal for 15 minutes. Pilot sessions involving 3-4 animals engaging in social interactions were also conducted. Animal behavior was manually labelled based on the video.

For comparison, similar behavioral recording were conducted using a tethered system (OpenEphys or Intan technologies digital acquisition systems). These are very widely used extracellular electrophysiology systems. In this case, mice carried a pre-amplifier connected to the implanted electrode and a thin tether (∼1.6 mm diameter) to bring the digitized signals to the amplifier board.

### Pupil tracking

Pupil dynamics in behaving mice were tracked using a head-mounted miniaturized camera. In the current implementation, we use a miniaturized CMOS camera with a resolution of 320 × 320 pixels and a minimum focal distance of 4 mm. We characterized the spatial resolution of the camera as a function of the distance between the camera and the eye, resulting in 26.3, 50, and 83.3 μm per pixel at distances of 4, 8, and 12 mm, respectively (Extended Data Fig. 5b). The temporal resolution of the camera recording was 16 Hz, which is sufficient to capture physiological pupil dynamics in mice^52^. In addition, the WILD platform can interface with a variety of other micro-cameras through standard industry interfaces such as the Serial Peripheral Interface (SPI), allowing higher-performance imaging modules to be integrated if needed.

Pupil diameter was automatically extracted using DeepLabCut (v3.0.0). Video frames were first manually labeled to train the network. Marker points were added to track both pupil and eye contour (Extended Data Fig. 5c). Pupil metrics were then normalized by eye width, as previously reported^53^.

Due to limited on-chip computational resources, complex image-processing libraries such as OpenCV are not currently supported for real-time pupil tracking on WILD.

### Outdoor recordings and analysis

Mice and rats’ outdoor recordings were conducted in enclosures at one of Cornell’s Field Stations. Enclosure dimensions ranged from 6 × 6 m^2^ to 15 × 38 m^2^. Walls were stainless steel panels inserted several inches deep into the ground. Enclosures were covered with netting to prevent birds and other predators from entering. Multiple video cameras and other types of sensors (e.g., radio-frequency receivers, ultra-wide band antennas) were placed in the enclosures to track animal movements. Food and water was exclusively provided in 9 wooded boxes (‘shelters’). Animal surgical and probe implantation procedures were identical to those of animals recorded inside the laboratory, but implants were encased in custom 3D-printed headstages. All male mice were implanted with WILD devices, which they carried for the duration of the experiment (in some cases for more than two weeks). Data included in this study were collected during free range exploration (between nights 1-15), enabling chronic monitoring of neural activity and behavior in naturalistic conditions. Electrodes were implanted in the dorsal hippocampus. After surgical implants, mice were allowed to recover during a week, after what they were transferred to the field station. All animals were dropped into the field enclosures the same day, at the same time and in the center of the paddock. Researchers monitored animals 24 / 7 and changed batteries daily. Animal position and identity were continuously tracked using a custom ultra-wideband (UWB) module attached to the main WILD board which position was triangulated through antennae placed around the enclosure, providing real-time localization (∼ 10 cm resolution) at ∼7-10 Hz. Position data were later down sampled to 4 Hz. For trajectory visualization, a speed filter (> 300 cm/s) was used to remove outliers, followed by smoothing based on median step size (kernel size = 3 samples). Trajectories were plotted as scatter plots on the aerial image of the enclosure.

Mice distance traveled was computed from UWB position data. Movement was defined as periods where instantaneous speed exceeded 5 cm/s and step distance was below 50 cm (to exclude tracking artifacts). Total distance per hour was summed and expressed in meters per hour (m / hr). Day and night classification was based on astronomical sunrise and sunset times for Ithaca, NY (42.44 N, 76.50 W), computed using the Astral library. Hour between sunset (∼19:00) and sunrise (∼6:00) were classified as *night*, the rest were classified as *day*. For day versus night comparisons, mean hourly distance was computed per animal for the day and night hours separately.

For place cell analysis, all putative hippocampal pyramidal cells with firing rates >= 0.1 Hz were included. Spatial rate maps were constructed by binning spiking data and occupancy into 3 cm spatial bins and smoothing with a Gaussian kernel (sigma = 3 bins). Only periods with animal speed > 2 cm/s and occupancy > 0.01 s per bin were included. Spatial information was computed in bits per spike^54^. A neuron was classified as a place cell if it met all of the following criteria: 1) spatial information significantly above chance (p < 0.05, shuffle test); 2) at least one quality-filtered place field (contour at 20% of local peak, minimum and maximum area 25 and 500 cm^2^ respectively, field firing rate > 1.5 mean firing rate; 3) peak firing rate >= 1.0 Hz and 4) total spike count >= 50. Place cells were further validated using a Poisson enrichment test with Benjamini-Hochberg FDR correction (q < 0.05) and leave-one-out robustness test (>= 5 runs, 100% pass fraction).

All quantifications for mice data from field recordings are from the same 9 animals. Statistical comparisons between day and night were performed using the two-sided Wilcoxon rank-sum test. Significant threshold was set at alpha = 0.05. Boxplots display the median, interquartile range (25^th^ - 75^th^ percentiles), and whiskers extending to 1.5 multiplied by interquartile range (IQR). Circadian plots display mean values +/- SEM across animals.

### 3D maze experiment

The 3D maze consisted in an elevated platform (46 cm diameter) with a movable ramp leading to a larger arena (1 m diameter). The platform had a single entrance to the empty space below it which features 50 small holes (5 mm diameter, 3 mm deep). The task of the mice was to find the shortest path to descend the platform, find the entrance below and, once inside, find a hidden water reward. Each trial ended once the animal obtained the reward, after which it was returned to its home cage. The relative position of the descending ramp and entrance, as well as the location of the hidden reward changed every session, forcing mice to learn a new efficient trajectory. In addition, the experiment was conducted in the darkness, so animals must rely on path integration and memory to solve it. The maze was thoroughly cleaned with 50% isopropyl alcohol between trials. Mice were implanted with silicon probes and optic fibers in the dorsal hippocampus. In some of the sessions, different optogenetic perturbations were applied (see next section).

### Closed-loop optogenetic manipulations with WILD

A two-channel optogenetic stimulation module was developed to interface with the WILD platform. The module is controlled by a microcontroller (STM32G071, STMicroelectronics), which regulates stimulation parameters and communicates with the WILD system via a 4-pin GPIO interface. A dual-channel current driver (TPS61151, Texas Instruments) provides independent control of two laser diodes or LEDs, with a maximum output current of 35 mA per channel. Stimulation currents are set by the microcontroller using onboard digital-to-analog converters. Key stimulation parameters, including intensity, duration, duty cycle, and waveform, are configured through commands from the WILD system. Arbitrary waveforms can be easily programmed. The optogenetic module measures 11.3 × 9.5 mm^2^ and weighs 0.6 g, including two mounted laser diodes (PL450B, Thorlabs).

WILD can be configured to deliver multiple types of optogenetic manipulations. Importantly, its advanced real-time signal processing capabilities allow using diverse types of signals, like neural activity or IMU, to condition on demand optogenetic stimulation (i.e., “closed-loop manipulations”). We illustrated two general case applications, such as the detection and disruption of a transient neural event (hippocampal sharp-wave ripples), and a phase-specific stimulation during hippocampal theta oscillations. Theta closed-loop and ripple closed-loop experiments were conducted on separate days, with control and experimental groups tested in an interleaved manner. Prior to the closed-loop experiments, the laser power was adjusted based on the amplitude of the evoked response. Theta closed-loop stimulation used a lower-power laser that did not evoke a clear neural response, whereas ripple closed-loop stimulation required a higher power sufficient to elicit an identifiable evoked response that interrupted ripple events.

### Software development

The embedded software was developed in C and compiled using the ARM toolchain. The PC control interface was implemented in C# with Microsoft Visual Studio 2022, providing functions for real-time device control and data download.

### On-board real-time signal processing with WILD

WILD embedded a signal processing pipeline and a TinyML inference engine. The signal processing pipeline consists of two parallel units for detecting biomarkers of LFPs. Data from the main ephys buffer is fed to the signal processing pipeline through an optional channel mixer that supports any linear combination of all recording channels. This allows for operations like noise subtraction, current-source-density, or extraction of principal component analysis (PCA) or independent component analysis (ICA). An immediate anti-aliasing filter is then used to low-pass filter the signal to 500Hz before downsampling to 1250Hz. A variety of first-order bandpass filters can be selected to filter the signal to the band of interest, and then the power envelope of the signal can be extracted. Two different envelop extraction techniques can be selected: 1) rectification-based: rectification of bandpassed signal, followed by lowpass filtering. 2) Hilbert transformer-based: A set of Hilbert transformer filters designed to introduce a −90° phase shift to the band-of-interest from previous bandpass filters, which will form an analytic signal together with the signal before HT. Hence, the phase angle can be obtained by calculating the arctangent of the ratio between the phase-shifted and original signals, while the envelope can be derived from the magnitude of these two signals combined. A threshold that is either calculated by multiplying a coefficient of the mean baseline level or arbitrarily set through wireless commands is used for event detection. Following event detection, a TinyML-based noise identification pipeline is triggered to reject artifacts using a 0.5s signal window preceding the event. A second TinyML engine performs behavior classification by continuously processing 1s segments of IMU data. The identified behavior labels are logged and transmitted wirelessly to the PC, enabling flexible automation of experimental control. TinyML models are pre-trained on a PC and compiled to firmware before experiments.

### Neural data analysis

Neural data analysis was conducted using MATLAB and Python. Electrophysiology data is processed with a custom MATLAB pipeline including LFP generation, noise channel rejection, automatic sleep scoring^55^ and spike sorting^56^, followed by manual curation in a Python GUI^57^. Single-unit activity was binned in 25-ms intervals, and neuronal assemblies were identified using ICA, with significance determined by null models, as implemented in a Python package (neuro_py^58^). Patterns were sign-aligned, and assembly activity was computed by projecting normalized spiking activity onto these patterns^59^. Spectra analysis of LFPs was performed with Gabor transformation^60^. SWR detection is performed as follows. Candidate SWRs were required to exceed 2 SD above the mean power envelope and have durations between 20 and 600 ms to be classified as SWRs. For Theta wave analysis, LFPs were first filtered with a zero-phase shifting Butterworth filter with a passband between 4 and 8 Hz, and the instantaneous phase was calculated by extracting the phase angle from the analytic signal produced by the Hilbert Transform. To compare the recording quality between tethered and WILD system. Same-day recordings were performed with a tethered system (RHD2000, Intan Technologies), followed by recording performed with WILD system. Recordings were segemented according to sleep states(Non-rapid eye movement, NREM; Rapid eye movement, REM) based on sleep scoring of hippocampus LFP. Continuous segments longer than 10 seconds were selected for analysis. Power spectral density (PSD) was computed using Welch’s method (512-point windows, no overlap, 625-point FFT). Median PSDs were calculated for each state, and distributions were statistically compared across devices using two-sample t-tests with Bonferroni correction. Spectral correlations were quantified using Pearson correlation of log-transformed PSDs. Noise performance was assessed by comparing recordings from grounded channels in both the tethered system and the WILD system. Signals were high-pass filtered with a second-order Butterworth filter (cutoff: 1 Hz), and RMS noise was calculated across channels. Noise levels were statistically compared using two-sample t-tests.

### Theta phase coherence

To quantify dynamic communication between the hippocampus and lateral septum, we measured time-resolved theta phase synchronization^61^. Specifically, we extracted the instantaneous phase of the 4-12 Hz theta band via the Hilbert transform and calculated the continuous Phase Locking Value (PLV)^62^ by averaging the complex phase differences over a 150 ms sliding window.

### Behavioral analysis

Over-the-head camera was used to record animal behaviors for indoor experiments. DeepLabCut (v3.0.0) was used for tracking animal positions with at least 3 track points(nose, body, tail). After position tracking, animal behaviors were manually inspected frame-by-frame and classified into motif, then cross-checked by two different experts, based on previously reported standards^63,64^. USVs were processed with the DeepSqueak toolbox^65^. USVs were first automatically detected, followed by manual curations to identify all vocalization events. USV features, including contour shape, duration, and frequency, were used for visualizing USV features with t-SNE^66^. Call types were clustered with an autoencoder and USV contour. 9-axis IMU data were processed with an AHRS filter from MATLAB. To compare the head movement of two socially interacting animals in tethered and untethered scenarios, a pair of animals with a tethered headstage with 3-axis accelerometers (RHD2000, Intan Technologies) or a WILD device. Principal component analysis was applied to accelerometer data to eliminate the device positioning differences among animals prior to pitch and roll calculation.

Mouse behaviors were manually labelled from inspection of recoded videos and categorized as social or non-social. Non-social behavior was defined as periods in which the animals were neither interacting nor directly facing one another. Social behaviors included nose sniffing (direct nose contact with the other animal’s nose or face), body sniffing (nose contact with the trunk of the other animal’s body), and anogenital sniffing (nose contact with the other animal’s anal/genital region). Additional social behaviors included chasing (one animal running away while the other follows directly behind), approaching (one animal actively moving towards another), and looking (both animals facing one another but without nose contact). Tail rattling and fighting were not observed during recording sessions.

### Behavior manifold embedding

Behavioral manifolds were constructed from synchronized pose and IMU features, following the general logic of prior wavelet-based unsupervised behavior-mapping approaches^67^. Body geometry was computed from tracked landmarks and combined with z-scored IMU roll, yaw, and pitch signals, with optional inclusion of inter-animal distance. These features were downsampled and transformed into 1-30 Hz time-frequency representations using continuous wavelet transforms. Wavelet amplitudes across channels and frequencies were concatenated and normalized at each time point, and pairwise Jensen-Shannon distances were computed between behavioral states. he resulting distance matrix was embedded into two dimensions with semi-supervised UMAP, in which behavioral labels were incorporated to modestly guide the organization of the manifold while preserving the structure imposed by the feature space.

### Onboard machine learning

Model trainings were performed in Python. Neural and behavioral data were loaded from electrophysiology (LFP) and IMU. Raw accelerometer signals were converted to physical units (g) and preprocessed to obtain pitch, roll, and derived features, including signal magnitude and jerk. Labels were loaded from manually curated labels based on DLC tracking. To address class imbalance, data were resampled by oversampling the minority classes to match the majority class. Labels were one-hot encoded for training. The dataset was split into training and validation sets (80/20). Model architectures were implemented in TensorFlow. The network was based on gated recurrent unit (GRU) layers and 1D Convolution Neural Networks (1D CNN), with hyperparameters optimized using grid search with cross-validation. Training was performed with categorical cross-entropy loss and Adam optimizer on a desktop PC with a GPU (Nvidia GTX 1080). The best-performing models were converted to TensorFlow Lite with post-training quantization for deployment on embedded hardware. Trained models were converted to Tensorflow-lite models, then compiled with STEdgeAI (STMicroelectronics). All analysis code was implemented in MATLAB 2020a and Python 3.9.

## Data availability

Data are available in a Zenodo repository (doi: 10.5281/zenodo.18879184). Due to their large size the original binary data files that support the findings of this study are available from the corresponding author upon request.

## Code availability

Instructions, design files, source code, and compiled binaries are available in a Git repository (https://github.com/ayalab1/Neurologger). Scripts used for data analysis are available on the same Zenodo repository (doi: 10.5281/zenodo.18879184). The device manual and operation instructions are available at https://ayalab1.github.io/Neurologger/. This code is available open-source under gpl-3.0 license.

## References

1. Steinmetz, N. A. et al. Neuropixels 2.0: A miniaturized high-density probe for stable, long-term brain recordings. Science 372, eabf4588 (2021).

2. Zong, W. et al. Large-scale two-photon calcium imaging in freely moving mice. Cell 185, 1240–1256.e30 (2022).

3. Pisano, F. et al. Depth-resolved fiber photometry with a single tapered optical fiber implant. Nat. Methods 16, 1185–1192 (2019).

4. Taal, A. J. et al. Optogenetic stimulation probes with single-neuron resolution based on organic LEDs monolithically integrated on CMOS. *Nat*. Electron. 6, 669–679 (2023).

5. Mathis, A. et al. DeepLabCut: markerless pose estimation of user-defined body parts with deep learning. Nat. Neurosci. 21, 1281–1289 (2018).

6. Weinreb, C. et al. Keypoint-MoSeq: parsing behavior by linking point tracking to pose dynamics. Nat. Methods 21, 1329–1339 (2024).

7. Nourizonoz, A. et al. EthoLoop: automated closed-loop neuroethology in naturalistic environments. Nat. Methods 17, 1052–1059 (2020).

8. Gomez-Marin, A., Paton, J. J., Kampff, A. R., Costa, R. M. & Mainen, Z. F. Big behavioral data: psychology, ethology and the foundations of neuroscience. Nat. Neurosci. 17, 1455–1462 (2014).

9. Kennedy, A. The what, how, and why of naturalistic behavior. Curr. Opin. Neurobiol. 74, 102549 (2022).

10. Tsoar, A., et al. Large-scale navigational map in a mammal. Proc. Natl. Acad. Sci. 108, E718–E724 (2011).

11. Takahashi, S., Hombe, T., Matsumoto, S., Ide, K. & Yoda, K. Head direction cells in a migratory bird prefer north. Sci. Adv. 8, eabl6848 (2022).

12. Kendall-Bar, J. M. et al. Brain activity of diving seals reveals short sleep cycles at depth. Science 380, 260–265 (2023).

13. Zhou, A. et al. A wireless and artefact-free 128-channel neuromodulation device for closed-loop stimulation and recording in non-human primates. *Nat*. Biomed. Eng. 3, 15–26 (2019).

14. Ginosar, G. et al. Locally ordered representation of 3D space in the entorhinal cortex. Nature 596, 404–409 (2021).

15. Forli, A., Fan, W., Qi, K. K. & Yartsev, M. M. Replay and representation dynamics in the hippocampus of freely flying bats. Nature 645, 974–980 (2025).

16. Jung, T. et al. A wireless subdural-contained brain–computer interface with 65,536 electrodes and 1,024 channels. *Nat*. Electron. 8, 1272–1288 (2025).

17. Lin, J., Zhu, L., Chen, W.-M., Wang, W.-C. & Han, S. Tiny Machine Learning: Progress and Futures [Feature]. IEEE Circuits Syst. Mag. 23, 8–34 (2023).

18. Guo, C. et al. Miniscope-LFOV: A large-field-of-view, single-cell-resolution, miniature microscope for wired and wire-free imaging of neural dynamics in freely behaving animals. Sci. Adv. 9, eadg3918 (2023).

19. Ide, K. & Takahashi, S. A Review of Neurologgers for Extracellular Recording of Neuronal Activity in the Brain of Freely Behaving Wild Animals. Micromachines 13, 1–11 (2022).

20. Yin, M. et al. Wireless neurosensor for full-spectrum electrophysiology recordings during free behavior. Neuron 84, 1170–1182 (2014).

21. Gagnon-Turcotte, G., Gagnon, L. L., Bilodeau, G. & Gosselin, B. Wireless brain computer interfaces enabling synchronized optogenetics and electrophysiology. in 2017 IEEE International Symposium on Circuits and Systems (ISCAS) 1–4 (2017). doi:10.1109/ISCAS.2017.8050345.

22. Ji, B. et al. Recent advances in wireless epicortical and intracortical neuronal recording systems. Sci. China Inf. Sci. 65, 140401 (2022).

23. Dennis, E. J. et al. Systems Neuroscience of Natural Behaviors in Rodents. J. Neurosci. 41, 911–919 (2021).

24. Zipple, M. N. et al. Competitive social feedback amplifies the role of early life contingency in male mice. Science 387, 81–85 (2025).

25. Egnor, S. R. & Seagraves, K. M. The contribution of ultrasonic vocalizations to mouse courtship. Curr. Opin. Neurobiol. 38, 1–5 (2016).

26. Ziobro, P., Woo, Y., He, Z. & Tschida, K. Midbrain neurons important for the production of mouse ultrasonic vocalizations are not required for distress calls. Curr. Biol. 34, 1107–1113.e3 (2024).

27. Chang, H. et al. Sleep microstructure organizes memory replay. Nature 637, 1161–1169 (2025).

28. Meyer, A. F., Poort, J., O’Keefe, J., Sahani, M. & Linden, J. F. A Head-Mounted Camera System Integrates Detailed Behavioral Monitoring with Multichannel Electrophysiology in Freely Moving Mice. Neuron 100, 46–60.e7 (2018).

29. Isaacson, M. et al. MouseGoggles: an immersive virtual reality headset for mouse neuroscience and behavior. Nat. Methods 22, 380–385 (2025).

30. Neunuebel, J. P., Taylor, A. L., Arthur, B. J. & Egnor, S. R. Female mice ultrasonically interact with males during courtship displays. eLife 4, e06203 (2015).

31. Oliva, A., Fernández-Ruiz, A., Leroy, F. & Siegelbaum, S. A. Hippocampal CA2 sharp-wave ripples reactivate and promote social memory. Nature 587, 264–269 (2020).

32. Buzsáki, G. Hippocampal sharp wave-ripple: A cognitive biomarker for episodic memory and planning. Hippocampus 25, 1073–1188 (2015).

33. Oliva, A., Fernández-Ruiz, A., Buzsáki, G. & Berényi, A. Role of Hippocampal CA2 Region in Triggering Sharp-Wave Ripples. Neuron 91, 1342–1355 (2016).

34. Fernández-Ruiz, A. et al. Long-duration hippocampal sharp wave ripples improve memory. Science 364, 1082–1086 (2019).

35. O’Keefe, J. & Dostrovsky, J. The hippocampus as a spatial map: Preliminary evidence from unit activity in the freely-moving rat. Brain Res. 34, 171–175 (1971).

36. Tang, J. C. Y. et al. Dynamic behaviour restructuring mediates dopamine-dependent credit assignment. Nature 626, 583–592 (2024).

37. Fernandez-Ruiz, A., Oliva, A. & Chang, H. High-resolution optogenetics in space and time. Trends Neurosci. 45, 854–864 (2022).

38. Ouyang, W. et al. An implantable device for wireless monitoring of diverse physio-behavioral characteristics in freely behaving small animals and interacting groups. Neuron 112, 1764–1777.e5 (2024).

39. Joo, H. R. & Frank, L. M. The hippocampal sharp wave–ripple in memory retrieval for immediate use and consolidation. Nat. Rev. Neurosci. 19, 744–757 (2018).

40. Girardeau, G., Benchenane, K., Wiener, S. I., Buzsáki, G. & Zugaro, M. B. Selective suppression of hippocampal ripples impairs spatial memory. Nat. Neurosci. 12, 1222–1223 (2009).

41. Robinson, H. L. et al. Large sharp-wave ripples promote hippocampo-cortical memory reactivation and consolidation during sleep. Neuron 114, 226–236.e6 (2026).

42. Khodagholy, D., Ferrero, J. J., Park, J., Zhao, Z. & Gelinas, J. N. Large-scale, closed-loop interrogation of neural circuits underlying cognition. Trends Neurosci. 45, 968–983 (2022).

43. Schomburg, E. W. et al. Theta Phase Segregation of Input-Specific Gamma Patterns in Entorhinal-Hippocampal Networks. Neuron 84, 470–485 (2014).

44. Newman, J. P. et al. ONIX: a unified open-source platform for multimodal neural recording and perturbation during naturalistic behavior. Nat. Methods 22, 187–192 (2025).

45. Marx, V. Neuroscientists go wireless. Nat. Methods 18, 1150–1154 (2021).

46. Park, S. I. et al. Soft, stretchable, fully implantable miniaturized optoelectronic systems for wireless optogenetics. Nat. Biotechnol. 33, 1280–1286 (2015).

47. Gutruf, P. et al. Fully implantable optoelectronic systems for battery-free, multimodal operation in neuroscience research. *Nat*. Electron. 1, 652–660 (2018).

48. Chen, J. C. et al. Self-rectifying magnetoelectric metamaterials for remote neural stimulation and motor function restoration. Nat. Mater. 23, 139–146 (2024).

49. Willmore, L., Cameron, C., Yang, J., Witten, I. B. & Falkner, A. L. Behavioural and dopaminergic signatures of resilience. Nature 611, 124–132 (2022).

50. Wang, X. et al. Empowering Edge Intelligence: A Comprehensive Survey on On-Device AI Models. ACM Comput Surv 57, 228:1–228:39 (2025).

## Methods-only references

51. Khodagholy, D. et al. Organic electronics for high-resolution electrocorticography of the human brain. Sci. Adv. 2, (2016).

52. Payne, H. L. & Raymond, J. L. Magnetic eye tracking in mice. eLife 6, e29222 (2017).

53. Chang, H. et al. Sleep microstructure organizes memory replay. Nature 637, 1161–1169 (2025).

54. Skaggs, W., McNaughton, B. & Gothard, K. An Information-Theoretic Approach to Deciphering the Hippocampal Code. in Advances in Neural Information Processing Systems vol. 5 (Morgan-Kaufmann, 1992).

55. Petersen, P. C., Siegle, J. H., Steinmetz, N. A., Mahallati, S. & Buzsáki, G. CellExplorer: A framework for visualizing and characterizing single neurons. Neuron 109, 3594–3608.e2 (2021).

56. Pachitariu, M., Steinmetz, N., Kadir, S., Carandini, M. & D H. K. Kilosort: realtime spike-sorting for extracellular electrophysiology with hundreds of channels. 061481 Preprint at 10.1101/061481 (2016).

57. Rossant, C., et al. phy: Interactive visualization and manual spike sorting of large-scale ephys data [Python]. The Cortical Processing Laboratory at UCL. (2023).

58. Harvey, R. ryanharvey1/neuro_py. (2025).

59. Harvey, R. E., Robinson, H. L., Liu, C., Oliva, A. & Fernandez-Ruiz, A. Hippocampo-cortical circuits for selective memory encoding, routing, and replay. Neuron 111, 2076–2090.e9 (2023).

60. Donoho, D., Maleki, A. & Shahram, M. Wavelab 850. Softw. Toolkit Time-Freq. Anal. (2006).

61. Sharif, F., Tayebi, B., Buzsáki, G., Royer, S. & Fernandez-Ruiz, A. Subcircuits of Deep and Superficial CA1 Place Cells Support Efficient Spatial Coding across Heterogeneous Environments. Neuron 109, 363–376.e6 (2021).

62. Lachaux, J.-P., Rodriguez, E., Martinerie, J. & Varela, F. J. Measuring phase synchrony in brain signals. Hum. Brain Mapp. 8, 194–208 (1999).

63. Gygax, M., Fortes, M. S., Voelkl, B., Würbel, H. & Novak, J. Rattling the cage: Behaviour and resource use of mice in laboratory and pet cages. Appl. Anim. Behav. Sci. 278, 106381 (2024).

64. Jabarin, R., Netser, S. & Wagner, S. Beyond the three-chamber test: toward a multimodal and objective assessment of social behavior in rodents. Mol. Autism 13, 41 (2022).

65. Coffey, K. R., Marx, R. E. & Neumaier, J. F. DeepSqueak: a deep learning-based system for detection and analysis of ultrasonic vocalizations. Neuropsychopharmacology 44, 859–868 (2019).

66. van der Maaten, L. & Hinton, G. Visualizing Data using t-SNE. J. Mach. Learn. Res. 9, 2579–2605 (2008).

67. Berman, G. J., Choi, D. M., Bialek, W. & Shaevitz, J. W. Mapping the stereotyped behaviour of freely moving fruit flies. J. R. Soc. Interface 11, 20140672 (2014).

