## Supplemental figures for "A wireless modular platform for neuro-behavioral recording and closed-loop manipulation in small animals"

### Extended Data and Figures

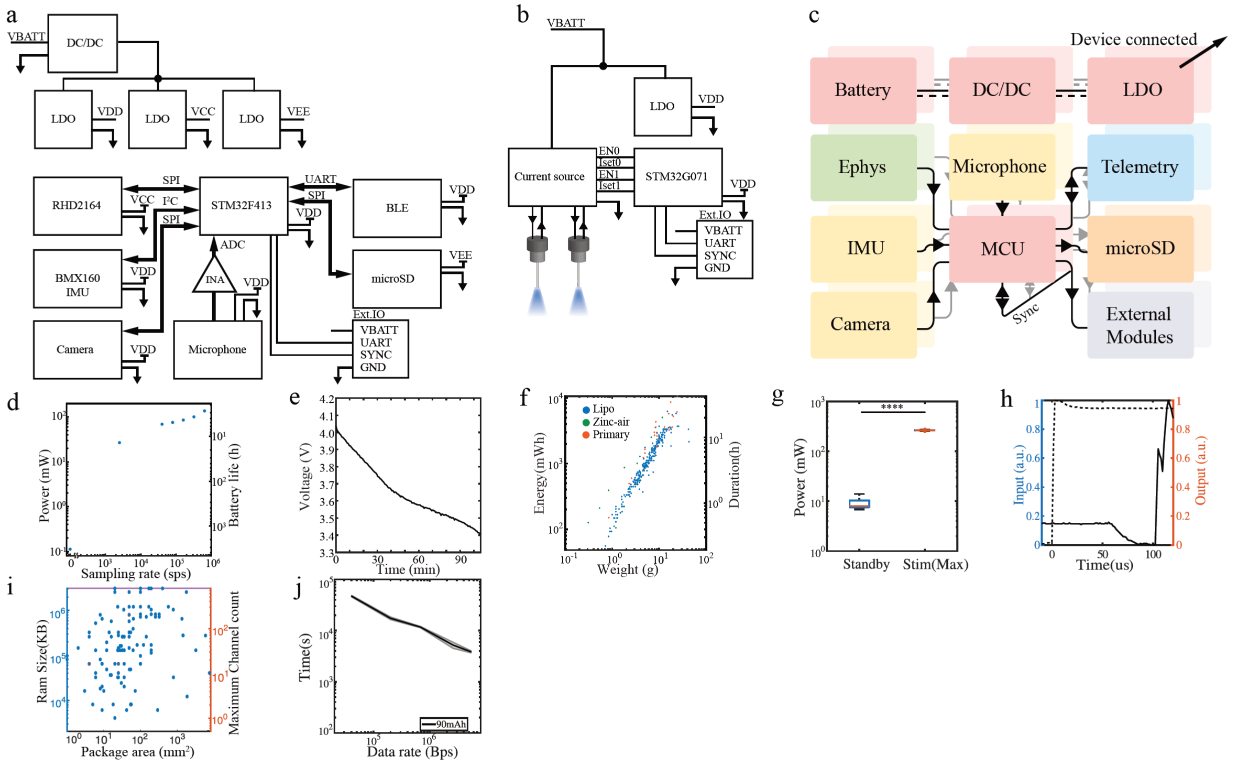

**Extended Data Figure 1: System schematics of WILD and performance metrics**

**a)** System diagram of WILD. The power system uses a high-efficiency DC-DC converter (TPSM83100, Texas Instruments) to operate across a wide input range (1.6-5.5 V), generating a 3.5 V rail. This is further regulated by three low-dropout regulators (TLV705, Texas Instruments), providing isolated 3.3 V supplies for the analog system (VCC), main digital system (VDD), and peripheral digital system (VEE). The core system integrates acquisition circuits, including a neural amplifier (RHD2164, Intan Technologies), 9-axis IMU (BMX160, Bosch Sensortec), camera (NanEye, ams-OSRAM), and ultrasonic microphone (SPH6611LR5H, Syntiant), followed by a 10× instrumentation amplifier (MAX4461, Analog Devices). Processing and communication are handled by a high-performance Cortex-M4 MCU (STM32F413, STMicroelectronics), a Bluetooth Low Energy subsystem (DA14580, Renesas), and a microSD card. External I/O (Ext. IO) carries data, synchronization signals, and power to connected subsystems. **b)** System diagram of the WILD optical stimulation module. A dual-channel current source (TPS61151, Texas Instruments) drives two laser diodes. A compact microcontroller (STM32G071, STMicroelectronics) controls the stimulation parameters, while a low-dropout regulator (LDO) supplies power to the MCU. **c)** Function diagram of WILD subsystems. **d)** Comparison of power consumption of the WILD system with different electrophysiology sampling rates (64 recording channels). **e)** Example battery discharge curve of WILD system powered by a 75 mAh lithium-polymer battery (64 recording channels at 20 kHz). **f)** Comparison of weight, energy of different batteries, and corresponding operation time based on high-speed recording mode (20 kHz, 64 channels) of WILD. **g)** Power consumption of the optogenetic stimulation module under standby conditions and during maximal stimulation (t-test $p<10^{-15}$). **h)** Time-resolved relationship between the stimulation input and emitted optical output, illustrating optogenetic stimulation module response dynamics and enabling measurement of power-on latency. **i)** Package area versus onboard RAM for candidate chips used in the logger design. Each point likely represents one chip. Smaller package area is better for a more compact and lightweight logger, whereas larger RAM size allows more buffering, preprocessing, or support for more recording channels. The right y axis (“minimal channel count”) converts RAM into an estimated number of channels. **j)** Estimated device operating time as a function of data rate. Log-scaled plot showing the tradeoff between communication bandwidth and battery life for a 90 mAh power source. Higher data rates are associated with shorter operating times, illustrating the power cost of increased data throughput.

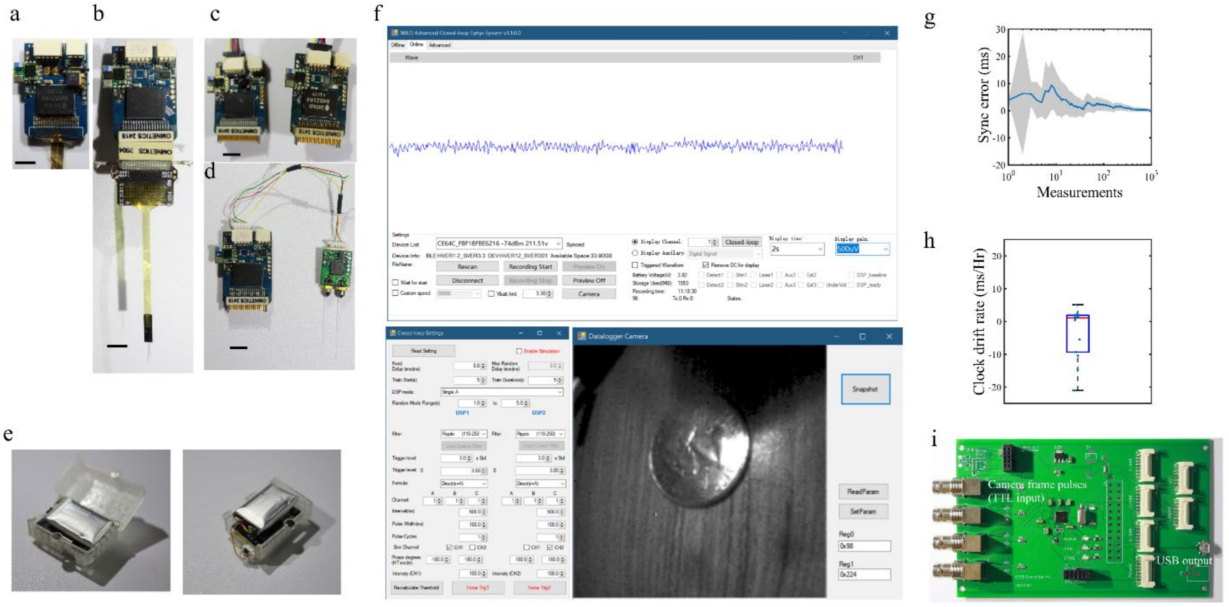

**Extended Data Figure 2: Wireless control of WILD with a custom framework**

**a)** WILD board mounted with paralyne-C flexible neural probe (Scale bar, 5cm). **b)** WILD board connected to a standard 64ch silicon probe (Cambridge Neurotech). Scale bar, 5cm. **(c)** Two WILD boards with standard Omnetics connectors synchronized through an external IO cable (Scale bar, 5cm). **d)** WILD board connected to optogenetic stimulator subsystem through a custom 4-wire ribbon (Scale bar, 5cm). **e)** Fully assembled device with custom designed 3D-printed case. Device is at the bottom, with battery on top. **f)** Graphical user interface of the WILD system on a Windows PC. The top panel shows neural activity (one channel) streaming in real time, and the main control menus for recording and visualization control. The bottom left panel shows real-time event detection and closed-loop stimulation controls. The right panel shows a video camera stream and controls. **g)** Clock synchronization error of WILD relative to the host PC during repeated over-the-air synchronization. The shaded area represents the 95% confidence interval. To maximize accuracy, the crystal oscillator was first calibrated by load tuning, followed by a time-of-flight-based estimation to align device time with the PC. Time estimation was then performed continuously throughout the recording to further refine synchronization. After this initial calibration, the device no longer required a continuous wireless link to the PC, allowing recordings from more animals than the typical seven-device connection limit. **h)** Measured clock drift with respect to the host PC. The box shows the interquartile range (IQR, 25th-75th percentile) with the median indicated by a central line. Whiskers extend to the most extreme data points within 1.5× IQR. Note that PC clock drift can only be corrected in steps of approximately 20 ms per day. **i)** Custom board to synchronize external devices, such as video cameras, with WILD. For a detailed list of operations enabled see Extended Data Table 2.

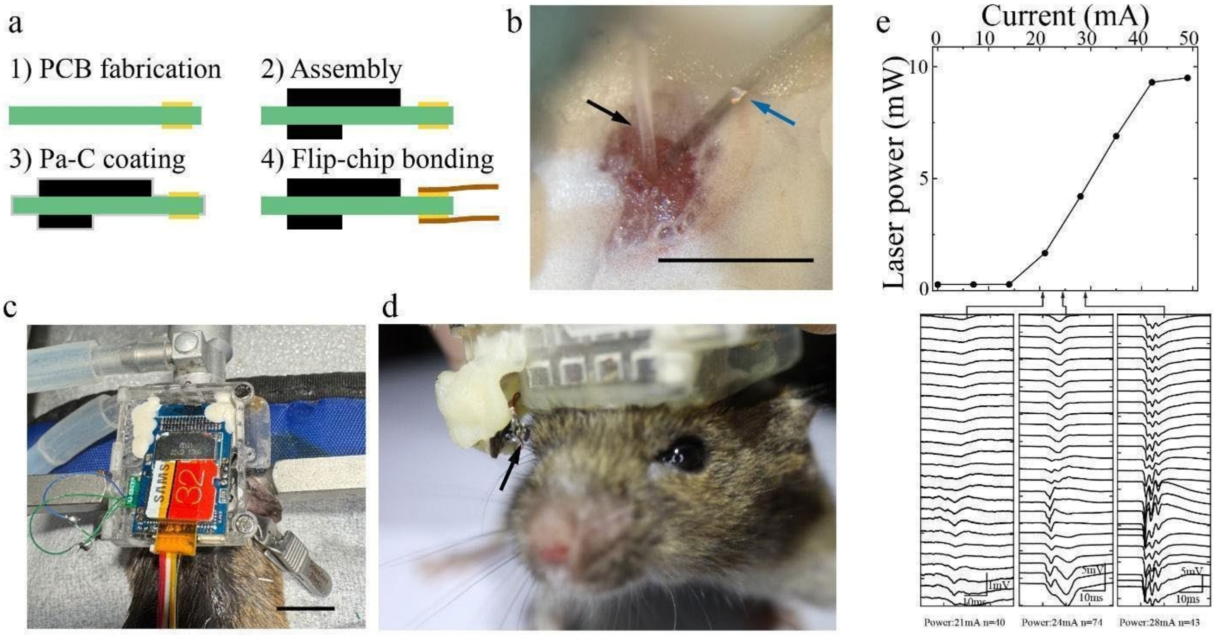

**Extended Data Figure 3: Integrated implantable device for closed-loop optogenetic experiments. a)** Illustration of the bonding process of weather-proof preparation of WILD. **b)** photo showing insertion of optical fiber (black arrow) and flexible neural probe (blue arrow) in the same implantation craniotomy (scale bar, 1mm) for simultaneous recording and optogenetic stimulation with WILD. **c)** Intraoperative image of an animal with an implanted device, electrodes, and encasing (bottom part only) in place (scale bar, 1cm). **d)** Photo of the pupillometry camera assembly in an implanted mouse. The arrow indicates the camera. **e)** WILD drives effective optogenetic stimulations. Upper panel: Blue light power measured from the tip of the optical fiber couple to the WILD optogenetic module as drive current increases. Bottom panel: Hippocampal evoked response as a result of optogenetic stimulation at increasing laser power.

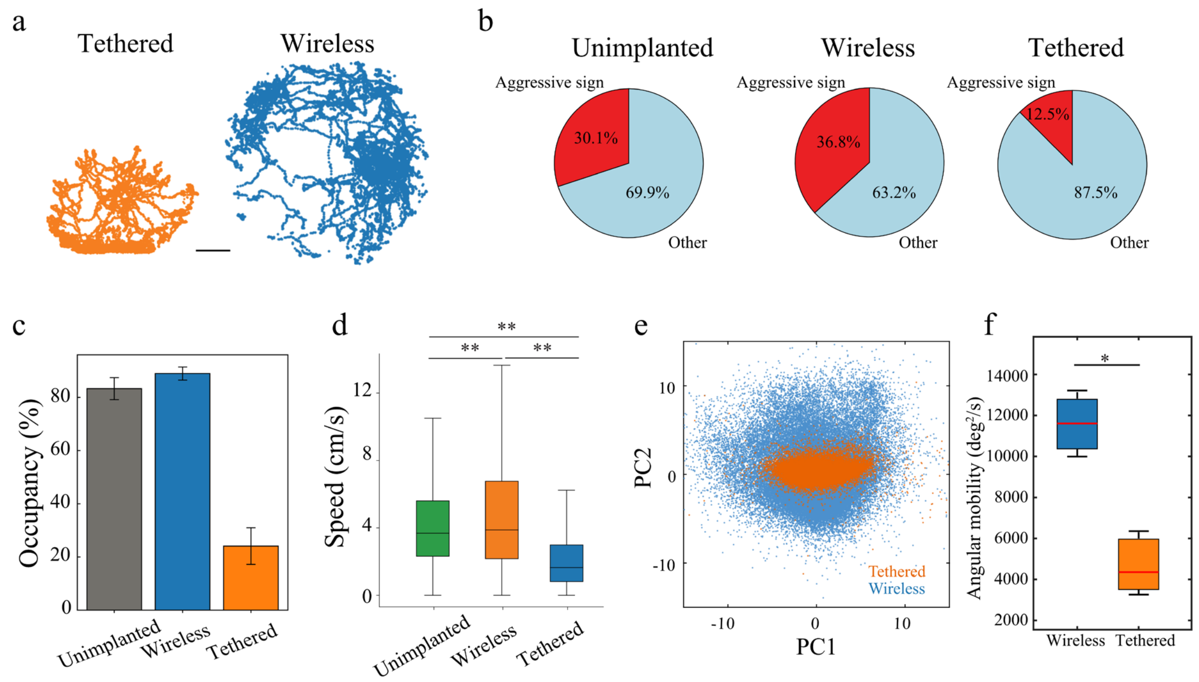

**Extended Data Figure 4: WILD enabled naturalistic behaviors in wireless recording settings**

**a)** Example tethered and wireless recorded mouse trajectories during a 15-minute open-field session (Scale bar, 5cm). **b)** Proportion of aggressive versus other social behaviors across experimental conditions. Each pie chart represents the composition of total interaction time within one condition: Control (unimplanted), Wireless, and Tethered male-male interaction sessions. Red slices denote time spent displaying aggressive signs (e.g., tail rattling, fighting, pushing), while light blue slices represent other non-aggressive behaviors. **c)** Wireless recording enhances spatial coverage during male-male social interaction. Bar plots show average occupancy coverage of unimplanted (gray), tethered (orange), and wireless (blue) recording conditions (mean ±1 SEM, n = 10 sessions; wireless vs tethered p = 0.0095, tethered vs unimplanted p = 0.0012, unimplanted vs wireless p = 0.57, Mann-Whitney U test). **d)** Moving speed distributions across different experimental conditions during social interaction. Boxplots show the distribution of linear speed (cm/s) for unimplanted, wireless, and tethered animals (** $p < 0.01$, Mann-Whitney U test, n = 10 sessions). **e)** Comparison of principal components of animal acceleration between wireless and tethered recordings. **f)** Comparison of angular mobility between wireless and tethered animals (N = 3). Statistical comparisons were performed using two-sided Kolmogorov-Smirnov tests (p= 0.032,*).

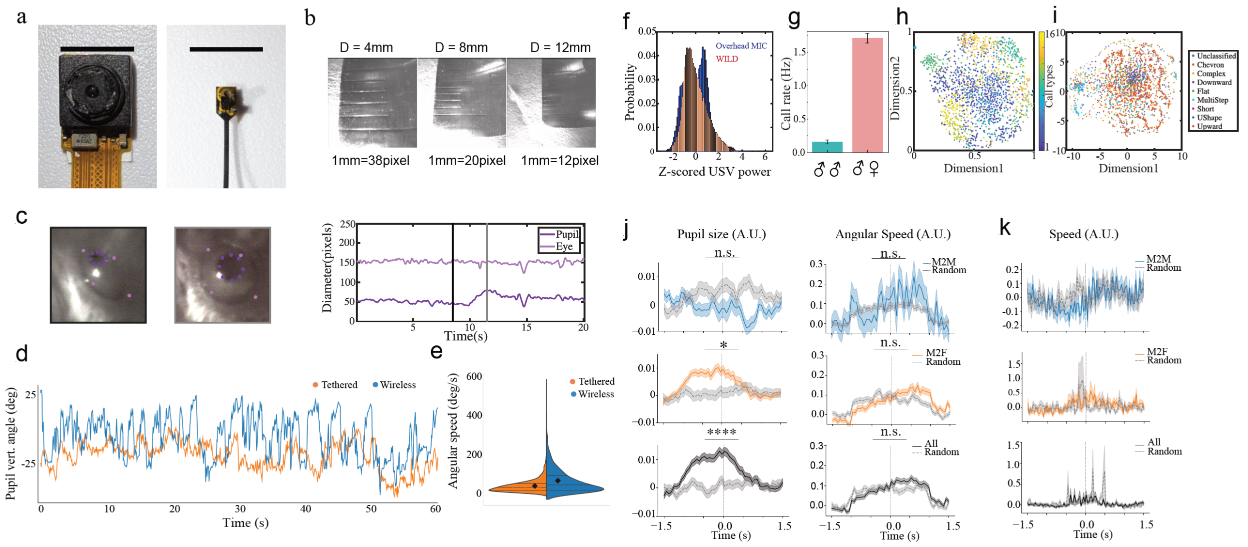

**Extended Data Figure 5: WILD enables pupillometry and USV recordings in behaving mice**

**a)** Size comparison between the Raspberry Pi camera used in previous studies[36] for tethered mice recordings and the new compact design. Scale bar, 10mm. **b)** Head-mounted camera performance at different working distances. Representative images acquired at object distances of 4, 8 and 12 millimeter, respectively, showing reduced effective spatial resolution at longer distances. **c)** Left, example of pupil and eye tracking with DeepLabCut; right, diameter of the eye (orange) and the pupil (blue) over a 20 s time window. **d)** Pupil vertical position over time. Blue trace: wireless; orange trace: tethered. **e)** Split violin plot showing the distribution of vertical angular speed in tethered (blue) versus wireless (orange) conditions. Each half violin represents the probability density of angular speed across 60 s of behavior, with black diamonds indicating the mean. The wireless method allowed for a broader and more variable range of eye movements compared to the more restricted distribution seen in the tethered condition (Mann-Whitney test: U= 362,246, p= $8.45 \times 10^{-16}$). **f)** comparison of USV power simultaneously detected by overhead external and head-mounted microphones. Bimodal distribution in overhead microphone denote USV power dependency animal location. **g)** Ultrasonic vocalization (USV) call rates during male-male and male-female interactions. Male-to-female interactions elicited significantly higher call rates compared to male-to-male interactions (t-test, $p < 0.0001$). Bars represent mean ± SEM. (male-to-male 155 calls from 3 sessions, male-to-female 3032 calls from 3 sessions) **h)** t-SNE plot comparing call signatures (spectra, duration) between different animal call types (N = 2,092). **i)** t-SNE plot of USV spectrograms classified using DeepSqueak[81]. Each point represents one vocalization color-coded by call type. **j)** Normalized pupil size and head angular speed aligned to male vocal calls, separated by interaction target. Top: Pupil size (A.U.) aligned to male-to-male (M2M, left) and male-to-female (M2F, right) calls. M2M: not significant ($p = 0.180$); M2F: $p < 0.05$ ($p = 0.024$). Bottom: Corresponding head angular speed (A.U.) around calls. M2M: not significant ($p = 0.880$); M2F: not significant ($p = 0.078$) (mean ±SEM; male-to-male 155 calls from 3 sessions, male-to-female 3032 calls from 3 sessions). **k)** Normalized head speed aligned to male vocal calls across different interaction contexts. Top: Male-to-male (M2M) interactions. Middle: Male-to-female (M2F) interactions. Solid colored lines show mean normalized speed around call onset (t = 0 s), with shaded areas indicating ±SEM. Dashed gray lines represent randomized control time points (mean ±SEM; $P > 0.05$ across all comparisons, paired t-test). Vertical dashed line marks call onset. (M2M: 155 calls from 3 sessions, M2F: 3032 calls from 3 sessions).

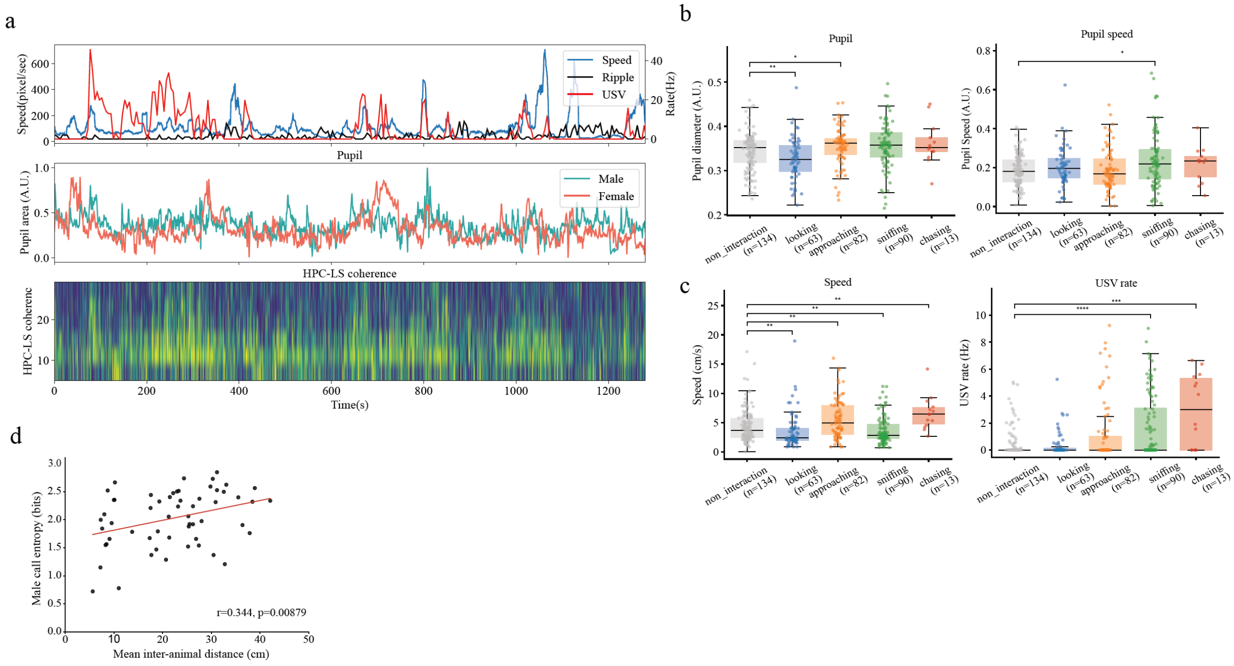

**Extended Data Figure 6: Physiological correlates of different social behaviors. a)** Top, example traces of ultrasound vocalization signals (USV, red), animal speed (speed, blue) and sharp-wave ripples (SWRs, black) rate. Middle, simultaneous recordings of pupil size in a male (blue) and a female (orange) during unrestricted interactions. Bottom, hippocampus-lateral septum (HPC-LS) spectral LFP during the same interaction window**. b)** Top, distribution of pupil diameter (left) and speed (right) during different social behaviors including looking, approaching, sniffing, chasing and no interaction. Note that pupil diameter significantly shrank and significantly dilated during looking (p=0.0065) and approaching(p=0.022) behaviors respectively compared to no interaction, while pupil speed was higher during sniffing compared to no interaction (p=0.013). Bottom, animal speed (left) and USV rate (right) during different behaviors. Note that animal's speed was significantly lower during looking(p=0.0026) and sniffing (p=0.0079), but significantly higher during approaching(p=0.0042) and chasing behaviors (p= 0.0010) respectively, compared to no interactions. USV rate was significantly higher during sniffing (p= $6.59 \times 10^{-5}$) and chasing (p= $1.6 \times 10^{-4}$) compared to no interactions. All tests are Mann-Whitney U tests **c)** Relationship between inter-animal distance (male and female) and entropy of the male emitted USV calls (n = 1683 calls over 20 s time windows) displayed a positive relationship, red line shows linear fit (r=0.3434, p=0.00879, n=57 time windows from 2 sessions). For each 20 s window, male call entropy was calculated as the Shannon entropy of the distribution of different call types and plotted against the corresponding mean inter-animal distance.

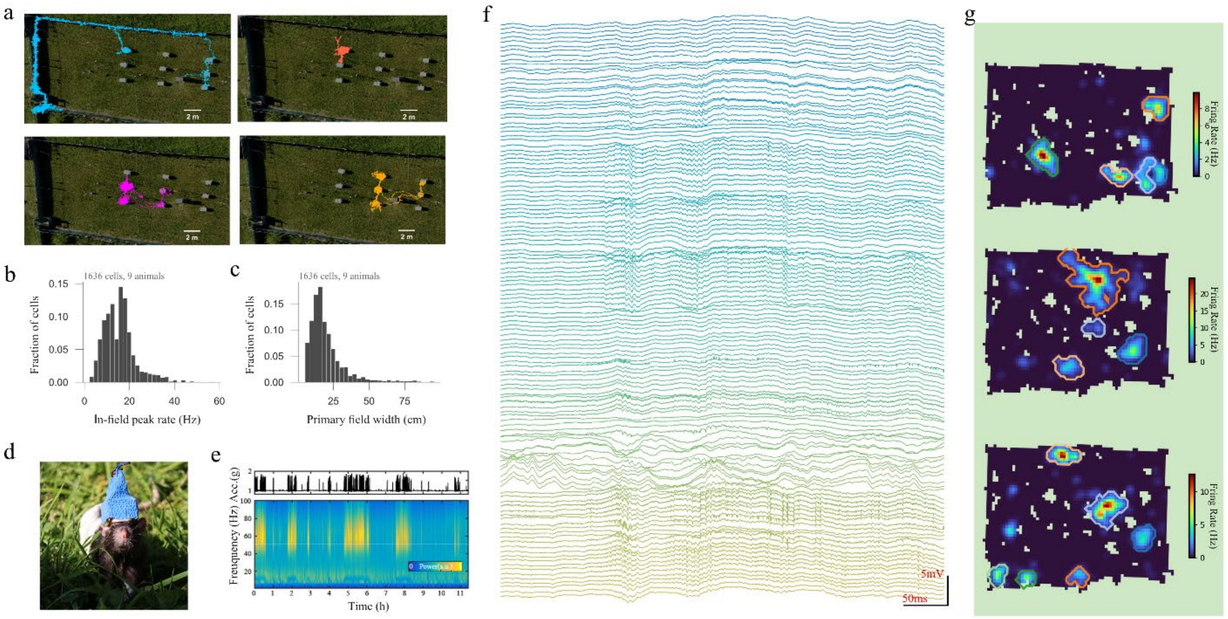

**Extended Data Figure 7: Individual animal trajectories and place cell population metrics. a)** Examples of individual UWB-tracked trajectories in outdoor enclosure for four simultaneously recorded mice on night 6, plotted over the aerial image of the field enclosure. Each panel shows a different animal (color coded as in Figure 5). **b)** Distribution of primary place field width (cm) for all identified place cells (n = 1636, n = 9 animals). **c)** Distribution of in-field peak firing rate for all identified place cells (n = 1636, n = 9 animals). **d)** Image of a rat with implanted WILD devices and silicon probes. **e)** Long duration (~12 hours) recording at low sampling rate (1250 Hz) of a rat implanted with WILD. **f)** Large-scale (128 channels) high sampling rate (20000 Hz) outdoor recording in a rat implanted with bilateral silicon probes and WILD. **g)** Examples of three dorsal hippocampal place cells from rats exploring an outdoor enclosure (2.4 x 3.4 m) obtained after recording with WILD.

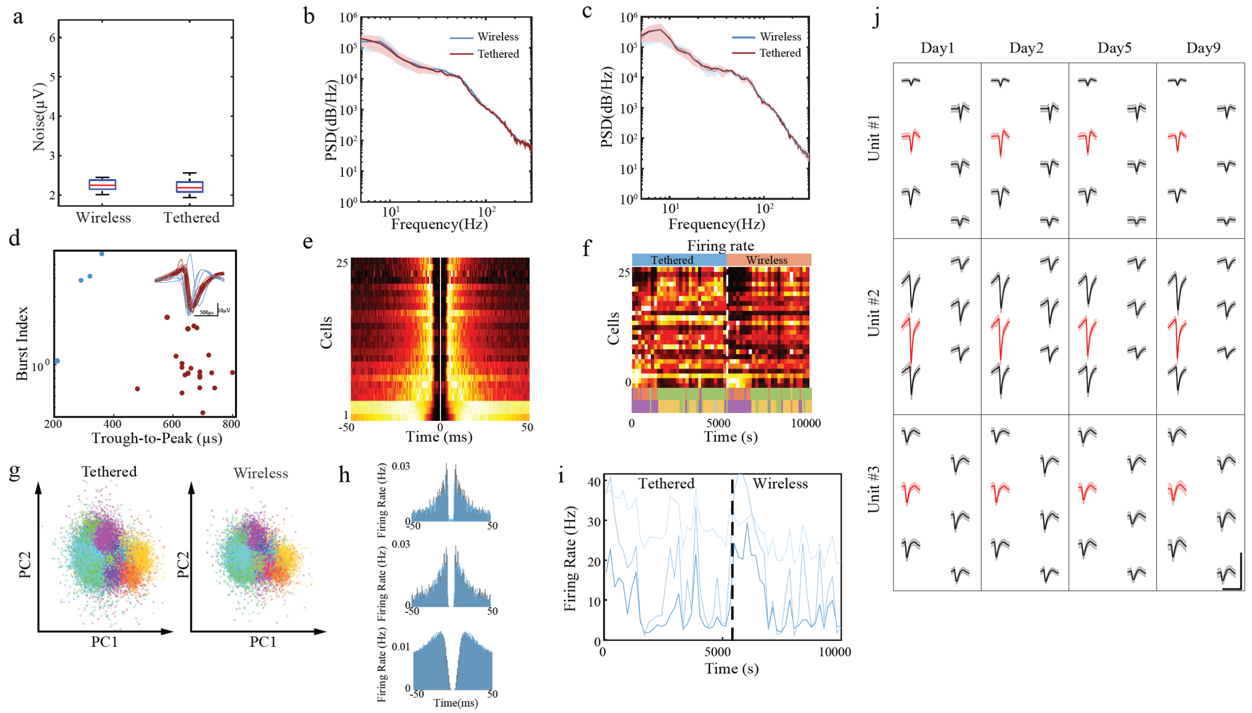

**Extended Data Figure 8: WILD enabled low-noise and high-quality single-unit electrophysiological recordings.** **a)** Comparison of root-mean-square noise of grounded channels between the WILD and Intan RHD2000 systems. $V_{WILD}$ = 2.92 ± 1.12 μV (N = 32) and $V_{Intan}$ = 2.80 ± 1.95 μV (N = 32); paired t-test, p = 0.757. **b)** Comparison of CA1 LFP power spectral density(PSD) between WILD and Intan RHD2000 system, during NREM sleep (N = 23). **c)** Comparison of CA1 LFP power spectral density (PSD) between WILD and Intan RHD2000 system, during REM sleep (N = 23). **d)** Comparison of burst index and trough-to-peak ratio of CA1 units (N = 25) recorded with flexible probes and WILD in behaving mice. **e)** Stacked ACGs of recorded CA1 units (N = 25). **f)** Firing rate of CA1 units vs time. The dashed line indicates the switch point from tethered recording to wireless recording**. g)** Comparison of the first and second principal components of spike waveforms from recorded units between tethered and wireless recordings. **h)** Selected ACGs of CA1 units during wireless (blue) and tethered (gray) recordings. **i)** Firing rates of selected units during tethered and wireless recordings. **j)** Examples of waveforms from stably recorded single units with a flexible neural probe over 9 days (Scale bar, 150 ms, 50 μV). Peak-amplitude channels are highlighted in red, and the shaded areas represent the standard error.

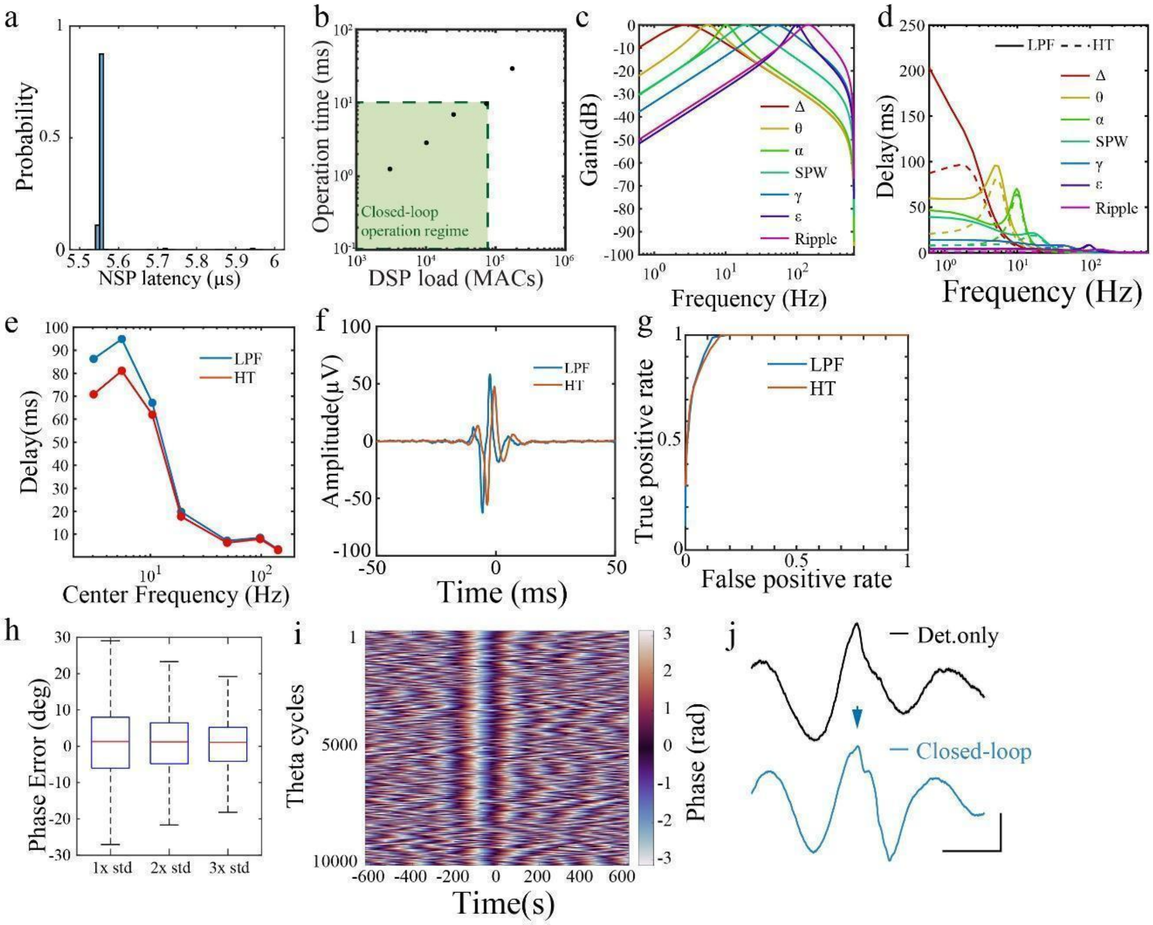

**Extended Data Figure 9: WILD embeds a fast, accurate online neural signal processor**

**a)** Signal processing latency of the embedded neural signal processor. Delay = 5.56 ± 0.09 μs, N = 387,561. **b)** WILD DSP operation time as a function of DSP load across different TinyML model sizes. The green box indicates the time requirement of closed-loop neural stimulation. **c)** Frequency responses of bandpass filters used in neural signature detection. Filters are first-order Butterworth filters with different passbands: Delta(Δ): 1-4Hz; Theta(θ): 4-8Hz; Alpha(α): 8-13Hz; Gamma(γ): 30-80Hz; Epsilon(ε): 80-120Hz; Sharp-wave(SPW): 12-30Hz; Ripple: 100-200Hz. **d)** Signal processing delays of NSP, showing Hilbert transformer-based NSP(HT, dashed line) have reduced delay compared to lowpass-filter-based NSP. **e)** Comparison between NSP delays based on LPF mode and HT mode across bandpass filters with different central frequencies. **f)** Trigger-averaged waveforms of CA1 ripple detections obtained using HT and LPF neural signal processing (NSP). **g)** ROC curves of ripple detection obtained using HT and LPF-based NSP, compared to a curated offline ripple detector. Area under curve: HT: 0.97218, N = 2,864; LPF: 0.97422, N = 2,075. **h)** Hippocampus CA1 theta phase decoding error of NSP phase decoder compared with offline decoder with Hilbert Transform, with signal amplitude in Theta band exceeding 1×/2×/3× standard deviation (STD, $N_{1\times STD}$ = 680,579; $N_{2\times STD}$ = 424,867; $N_{3\times STD}$ = 161,774). **i)** Trigger-averaged theta phase of phase-based NSP detections (phase window set to -15° to 15°, N = 10,085). **j)** Triggered average of LFPs for onboard-detected events during detection-only (N = 1,670) and closed-loop theta-wave intervention (N = 6,052) conditions (scale bar: 100ms, 100μV). Stimulation onset is indicated by the blue arrow.

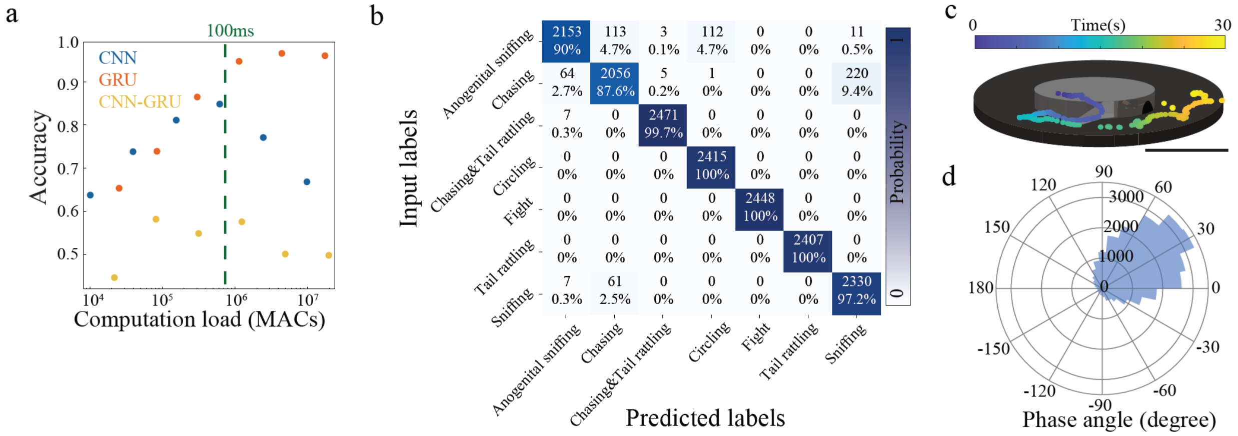

**Extended Data Figure 10: TinyML allows rapid computations for online processing during behavior.** **a)** Inference accuracy of different TinyML models on behavior identification compared with model size and model structures. Target inference times are indicated by a dashed line. **b)** Confusion matrix comparing onboard-predicted behavior labels with ground-truth reference labels (Behavioral events were annotated and quantified, including sniffing (n = 194), anogenital sniffing (n = 19), chasing (n = 108), chasing & anogenital sniffing (n = 3), chasing & tail rattling (n = 11), circling (n = 7), fighting (n = 6), pushing (n = 4), and tail rattling (n = 56)).**c)** Example of mouse trajectory during exploration of double-layer maze (scale bar: 50 cm). **d)** Histogram of real-time stimulation theta phases (N=15,728). Target was 30-45°.

| Name | Power (mW) | Channels | Sps | Mode | Weight (g) | Video | Audio | Motion sensor | Processing |
|---|---|---|---|---|---|---|---|---|---|
| This work (WILD) | 0.11 | 0 | 0 | Sleep | 1.48 | | | | |
| | 24.42 | 8 | $4\times10^4$ | Ephys, IMU | | | | Accelerometer/ Gyroscope/ Magnetosensor | Spectral power, tinyML |
| | 66.28 | 64 | $2\times10^5$ | Ephys, IMU | | | | | |
| | 101.45 | 64 | $6.8\times10^5$ | Ephys, IMU | | | | | |
| | 152 | 64 | $1.32\times10^6$ | Ephys, IMU | | | | | |
| | 194.04 | 64 | $2.6\times10^6$ | Ephys, IMU | | | | | |
| | 108.29 | 64 | $5.2\times10^5$ | Ephys, IMU, Audio | | | | | |
| | 143.01 | 64 | $1\times10^6$ | Ephys, IMU, Audio | | | 160KHz | | |
| | 171.56 | 64 | $1.64\times10^6$ | Ephys, IMU, Audio | | | | | |
| | 208.67 | 64 | $2.92\times10^6$ | Ephys, IMU, Audio | | | | | |
| | 180.53 | 64 | $3.08\times10^6$ | Ephys, IMU, Audio, Video | | 320 x 320 pixel, 16Hz | | | |
| | 182.94 | 64 | $3.56\times10^6$ | Ephys, IMU, Audio, Video | | | | | |
| | 204.81 | 64 | $4.2\times10^6$ | Ephys, IMU, Audio, Video | | | | | |
| | 245.28 | 64 | $5.48\times10^6$ | Ephys, IMU, Audio, Video | | | | | |
| Neuropixel Datalogger | 608 | 400 | $1.51\times10^7$ | Ephys, IMU | 15 | | | Accelerometer/ Gyroscope | |
| miniLogger32 | 335.4 | 32 | $9.6\times10^5$ | Ephys, IMU | 3 | | | Accelerometer/ Gyroscope | |
| Neurologger3 | 148 | 64 | $1.33\times10^6$ | Ephys, IMU, Audio | 1.96 | | 125KHz | Accelerometer/ Gyroscope/ Magnetosensor | |
| Neurologger2 | 121 | 4 | $3.84\times10^4$ | Ephys, IMU, Audio | 1.71 | | 200KHz | Accelerometer | |
| MouseLog-16B | 129.5 | 16 | $5\times10^5$ | Ephys | 1.9 | | | | |
| RatLog-32 | 229.4 | 32 | $1.02\times10^6$ | Ephys, IMU, Audio | 3.3 | | 200KHz | Accelerometer/ Gyroscope/ Magnetosensor | |
| RatLog-64 | 314.5 | 64 | $2.05\times10^6$ | Ephys, IMU, Audio | 4.92 | | 200KHz | Accelerometer/ Gyroscope/ Magnetosensor | |
| RatLog128 | 314.5 | 128 | $4.1\times10^6$ | Ephys, IMU | 7.92 | | | | |
| Gosselin, 2017 | 175 | 32 | $6.4\times10^5$ | Ephys, IMU | 2.8 | | | | |
| Muller, 2019 | 172 | 128 | $1.28\times10^5$ | Ephys, IMU | 7.4 | | | Accelerometer/ Gyroscope | Spectral power |
| Yin, 2014 | 145 | 96 | $1.92\times10^6$ | Ephys, IMU | 3.6 | | | Accelerometer | |
| CerePlex Exilis | 592 | 96 | $2.35\times10^6$ | Ephys, IMU | 9.87 | | | | |

| | | | | | |
|---|---|---|---|---|---|
| HD32 | 129.5 | 32 | $4\times10^4$ | Ephys | -- |
| HS-W | --- | 64 | $1.28\times10^6$ | Ephys | 2.59 |

**Extended Data Table 1: Device specifications.** Comparison of WILD specifications with those of previous wireless neural recording devices. Sources are academic publications of datasheets of commercially available devices.

| Cmd ID | Length (Bytes) | Function |
|---|---|---|
| 0x01 | 513 | Set recording parameter |
| 0x02 | 513 | Set DSP1 parameter |
| 0x03 | 513 | Set DSP2 parameter |
| 0x13 | 4 | Set stimulation intensity |
| 0x14 | 6 | Set detection threshold |
| 0x15 | 6 | Set detection level |
| 0x20 | 21 | Set stimulator1 parameter |
| 0x21 | 21 | Set stimulator2 parameter |
| 0x30 | 1 | Start recording |
| 0x31 | 1 | Stop recording |
| 0x40 | 1 | Start online previewing |
| 0x41 | 1 | Stop online previewing |
| 0x42 | 2 | Set preview channel |
| 0x51 | 1 | Start impedance testing |
| 0x60 | 2 | Manual stimulation trigger |
| 0x61 | 3 | GPIO control |
| 0x80 | 1 | Connection check |
| 0x82 | 4 | 2-way time delay estimation |
| 0x8A | 20 | Set device time |
| 0x90 | 1 | Read recording parameters |
| 0x91 | 1 | Read DSP1 parameter |
| 0x92 | 1 | Read DSP2 parameter |
| 0x9E | 1 | Camera snapshot |
| 0xA0 | 1 | Request deep sleep |
| 0xAB | 2 | Forced device reset |

**Extended Data Table 2: Bluetooth Command List.** List of operations that can be deployed in WILD from host computer via Bluetooth communication
